# A binary effector module secreted by a type VI secretion system

**DOI:** 10.1101/2021.07.29.453783

**Authors:** Yasmin Dar, Biswanath Jana, Eran Bosis, Dor Salomon

## Abstract

Gram-negative bacteria use type VI secretion systems (T6SSs) to deliver toxic effector proteins into neighboring cells. Cargo effectors are secreted by binding non-covalently to the T6SS apparatus. Occasionally, effector secretion is assisted by an adaptor protein, although the adaptor itself is not secreted. Here, we report a new T6SS secretion mechanism, in which an effector and a co-effector are secreted together. Specifically, we identified a novel periplasm-targeting effector that is secreted together with its co-effector, which contains a MIX (marker for type sIX effector) domain previously reported only in polymorphic toxins. The effector and co-effector directly interact, and they are dependent on each other for secretion. We termed this new secretion mechanism “a binary effector module”, and we show that it is widely distributed in marine bacteria.

## INTRODUCTION

One of the most diverse bacterial toxin delivery systems is the type VI secretion system (T6SS); it targets toxins, termed effectors, into either bacteria or eukaryotic neighboring cells in a contact-dependent manner (Pukatzki *et al*, 2006; Mougous *et al*, 2006; Hood *et al*, 2010; Pukatzki *et al*, 2007). Effectors possessing antibacterial activities are encoded together with a cognate immunity protein that prevents self-intoxication by physically binding the effector and antagonizing its activity at its subcellular destination (i.e., in the cytoplasm, membrane, or periplasm) (Russell *et al*, 2012, 2011).

T6SS effectors are loaded onto a secreted tail tube composed of stacked hexameric rings of Hcp proteins, which are capped by a spike complex comprising a VgrG trimer and a PAAR repeat-containing protein (hereafter, referred to as PAAR) that sharpens the tip of this structure (Nazarov *et al*, 2017; Wang *et al*, 2017; Shneider *et al*, 2013). The tail tube is propelled out of the cell by a contracting sheath structure that engulfs it inside the secreting bacterium (Basler *et al*, 2012). Effectors are deployed once the tail tube has penetrated a recipient cell.

Several mechanisms mediating the translocation of T6SS effectors into recipient cells have been characterized. The first characterized mechanism was the delivery of specialized effectors (also known as ‘evolved effectors’) (Pukatzki *et al*, 2007), a term referring to the three secreted tail tube components of the T6SS (Hcp, VgrG, and PAAR) when they are fused to a C-terminal toxin domain (Ma *et al*, 2017; Pukatzki *et al*, 2007; Shneider *et al*, 2013). A second type of effectors, known as cargo effectors, are toxin domain-containing proteins that non-covalently attach to one of the three secreted tail tube components (Bondage *et al*, 2016; Flaugnatti *et al*, 2016; Jana *et al*, 2019; Wettstadt *et al*, 2019; Hachani *et al*, 2014; Flaugnatti *et al*, 2020).

Many cargo effectors and PAAR-containing specialized effectors require cognate adaptor proteins. Adaptors function as chaperones that bind the effector and contribute to its stability and loading onto the T6SS tail tube. Four adaptor domains (DUF4123, DUF1795, DUF2169, and DUF2875) have been experimentally validated (Unterweger *et al*, 2015; Liang *et al*, 2015; Alcoforado Diniz & Coulthurst, 2015; Cianfanelli *et al*, 2016; Quentin *et al*, 2018; Bondage *et al*, 2016; Ahmad *et al*, 2020; Berni *et al*, 2019); co-adaptors have also been reported to occasionally participate in this process (Burkinshaw *et al*, 2018). Moreover, Hcp serves as a chaperone for several effectors that are loaded inside the Hcp tube (Silverman *et al*, 2013). Adaptors are commonly encoded adjacent to the effector, although there have been reports of adaptors encoded at a distant genetic locus (Ahmad *et al*, 2020). Importantly, the adaptors are not secreted, and the mechanism ensuring their intracellular retention and their dissociation from the effector remains unclear.

Members of the *vibrionaceae* family are Gram-negative bacteria prevalent in aquatic ecosystems (Boyd *et al*, 2015); they include established and emerging pathogens of humans and marine animals (Horseman *et al*, 2013). Many vibrios harbor at least one T6SS in their genome (Dar *et al*, 2018). These T6SSs are employed in interbacterial competition, anti-eukaryotic toxicity (virulence or antagonizing predation), or both (Salomon *et al*, 2013; Ray *et al*, 2017; Pukatzki *et al*, 2006; MacIntyre *et al*, 2010; Salomon *et al*, 2015; Hubert & Michell, 2020; Speare *et al*, 2018). *Vibrio parahaemolyticus*, a widespread emerging pathogen, is a major cause of seafood-borne gastroenteritis (Newton *et al*, 2012; Zhang & Orth, 2013) and of acute hepatopancreatic necrosis disease (AHPND) in shrimp (Tran *et al*, 2013; Lai *et al*, 2015). Pathogenic isolates of this bacterium encode a T6SS, termed T6SS1 (Li *et al*, 2017; Salomon *et al*, 2013; Yu *et al*, 2012), whose closely homologous systems are widespread in vibrios and other marine bacteria (Dar *et al*, 2018; Salomon *et al*, 2015; Ray *et al*, 2017). The activities and effector repertoires of this T6SS have been investigated in several *Vibrio* strains. Notably, in all four *V. parahaemolyticus* isolates in which T6SS1 has been experimentally investigated, as well as in the investigated homologous T6SSs in *V. alginolyticus* 12G01 and in *V. proteolyticus* NBRC 13287, a tricistronic operon is found at the beginning of the T6SS cluster (Salomon *et al*, 2014a; Ray *et al*, 2017; Salomon *et al*, 2015; Jana *et al*, 2019; Fridman *et al*, 2020). This tricistronic operon, corresponding to *vp1388-vp1390* in the *V. parahaemolyticus* type strain RIMD 2210633 (Fig. 1A), was implicated in interbacterial competition (Salomon *et al*, 2014a). Interestingly, both VP1388 and VP1390, as well as their *V. alginolyticus* and *V. proteolyticus* homologs, are secreted in a T6SS1-dependent manner (Ray *et al*, 2017; Salomon *et al*, 2015, 2014a). VP1389, encoded by the middle gene of the tricistronic operon, and its homologs contain an N-terminal signal peptide for periplasmic localization (Fig. 1A).

**Figure 1.**
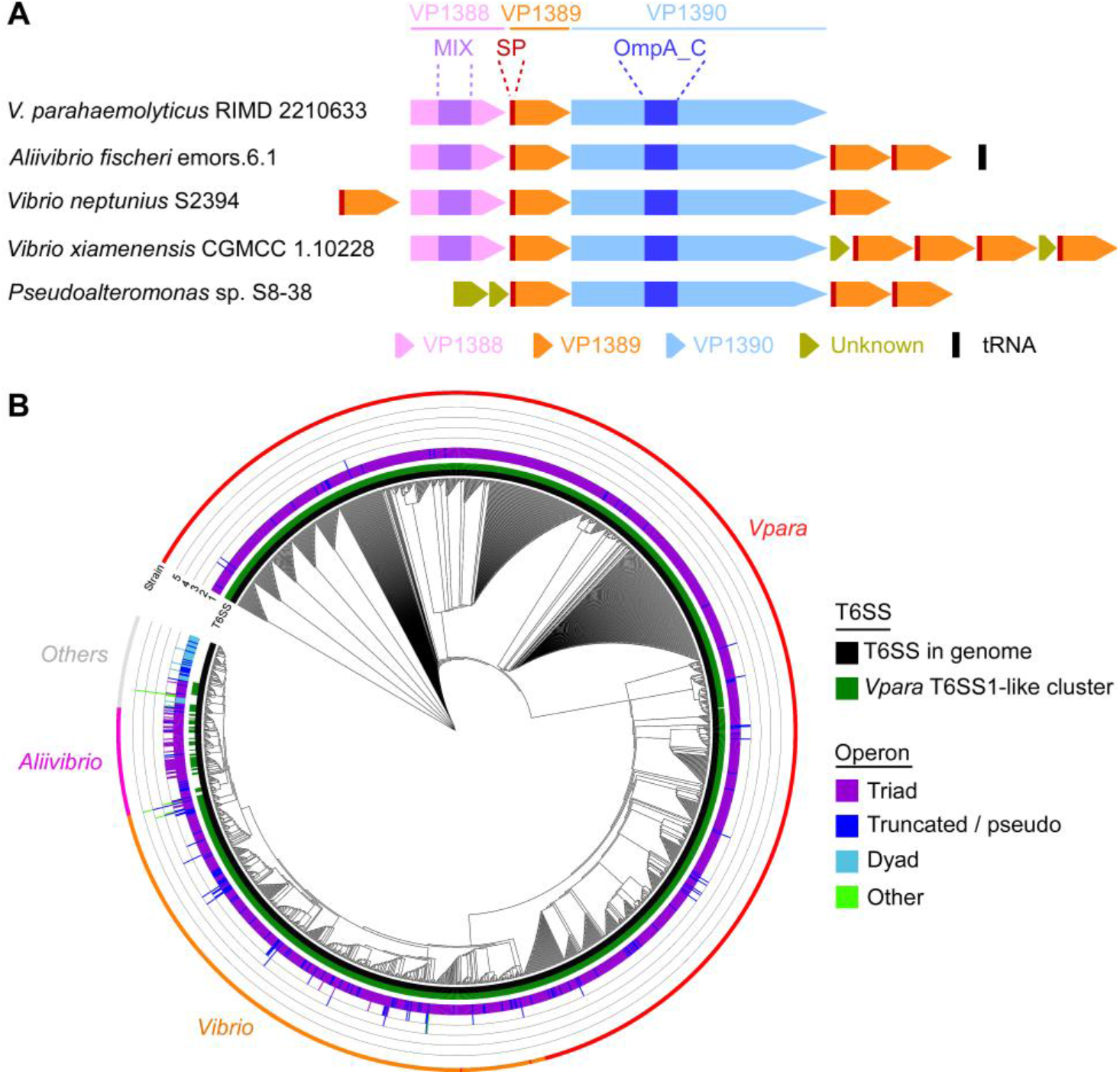
*vp1388-90* homologous operons are widespread in T6SS-encoding marine bacteria. **A)** Selected examples of the genetic structure of *vp1388-90* homologous operons. SP, signal peptide; MIX, Marker for type sIX effector. **B)** Distribution of *vp1388-90* homologous operons in bacteria. A phylogenetic tree of bacteria encoding homologous operons, based on the DNA sequences of *rpoB*. The presence or absence of T6SS in each genome is denoted in the inner rings (black and dark green). Intermediate bars indicate the number of complete (triad) and partial (dyad or truncated) homologous operons identified in each genome. An external ring denotes the group to which the bacterial strains belong. *V. parahaemolyticus* (*Vpara*) were annotated separately (red), as were *Aliivibrios* (pink).

Previously, we proposed that VP1388, containing a MIX (Marker for type sIX effector) domain that indicates a secreted T6SS substrate, is a T6SS effector and that VP1389 is an immunity protein (Salomon *et al*, 2014a); deletion of both genes render a prey strain sensitive to T6SS1-mediated attacks by a parental competitor, for which VP1388 is required. Exogenous expression of VP1389 in the prey restores immunity. However, the role of the third operon-encoded protein, VP1390, and the antibacterial activity mediated by this operon have remained unknown.

Here, we investigated the roles and activities of the proteins encoded by this T6SS-associated operon. Importantly, we found that both VP1388 and VP1390 are required for the antibacterial activity mediated by the tricistronic operon, and that their secretion is co-dependent. Furthermore, we demonstrated that VP1388 and VP1390 interact directly and are loaded together on the T6SS tail tube. Lastly, we revealed that VP1390, rather than VP1388, mediates antibacterial toxicity in the periplasm; its activity resulted in distinct morphological changes that led to cell lysis. We propose that VP1390 is a newly identified antibacterial T6SS effector that uses a novel secretion mechanism, whereby the MIX domain-containing VP1388 serves as its secreted co-effector.

## RESULTS

### Homologous operons of *vp1388-vp1390* are widespread in marine bacteria

Prior to characterizing the functions of the three operon-encoded proteins, VP1388, VP1389, and VP1390, we first set out to determine the operon’s distribution and conservation. To this end, we identified homologs of these three proteins in available bacterial genomes and investigated their genomic neighborhoods. Operons that encode homologs of all three proteins (hereafter, referred to as triads) were found in genomes of 1375 marine bacterial strains harboring T6SS, mostly belonging to the *vibrionaceae* family (Fig. 1 and Supplementary Datasets S1 and S2). In some genomes (e.g., *Aliivibrio*), more than one copy of the operon was detected. Often, multiple copies of the putative immunity protein, homologous to VP1389, were present within the operon or flanking it (Fig. 1A and Supplementary Dataset S2). These additional copies may represent orthologs that have been acquired via horizontal gene transfer or that have evolved to protect against non-kin toxins, since they often bear more sequence similarity to proteins encoded by other bacterial strains than to their neighbors. Notably, ~65% of the homologous operons were found in proximity to T6SS core proteins, usually at the edges of T6SS gene clusters (Supplementary Dataset S3), indicating their association with this secretion system.

Interestingly, homologs of VP1388 were almost exclusively found in triads. When an operon was truncated at the end of a contig or it included pseudogenes at the edge, thus hampering our ability to confidently determine the genetic composition of the operon, it was denoted as “Truncated/pseudo” (Fig. 1B). Nevertheless, a handful of instances in which VP1388 was found alone or only with a VP1390 homolog were detected (denoted as “Others” in Fig. 1B). Interestingly, we also found various genomes in which a VP1388 homolog is absent (e.g., in *Pseudoalteromonas*, *Bermanella*, and *Desulfoluna*); however, VP1390 and VP1389 homologs are present (denoted as “Dyads”; Fig. 1B). This observation suggests a link between VP1390 and VP1389. Remarkably, genomes encoding dyads did not encode a T6SS that is similar to *V. parahaemolyticus* T6SS1, as opposed to the vast majority of genomes encoding a triad.

### Only VP1389 is required for immunity against T6SS1-mediated toxicity

In a previous work, we showed that VP1389 was required for immunity against T6SS1-mediated aggression (Salomon *et al*, 2014a). However, we did not directly investigate whether VP1388 and VP1390 play a role in immunity. To test this, we deleted each of the three genes, *vp1388*, *vp1389*, and *vp1390*, individually and determined the ability of each mutant to defy intoxication by a wild-type attacker during competition. As shown in Fig. 2A, only *vp1389* was necessary for immunity against a T6SS1-mediated attack, whereas neither *vp1388* nor *vp1390* was required. Notably, deletion of *vp1388* resulted in slightly lower prey growth; however, this was not due to T6SS1-mediated toxicity of the attacker (Supplementary Fig. S1). These results indicate that the two secreted proteins, VP1388 and VP1390, do not play a role in immunity against T6SS1.

**Figure 2.**
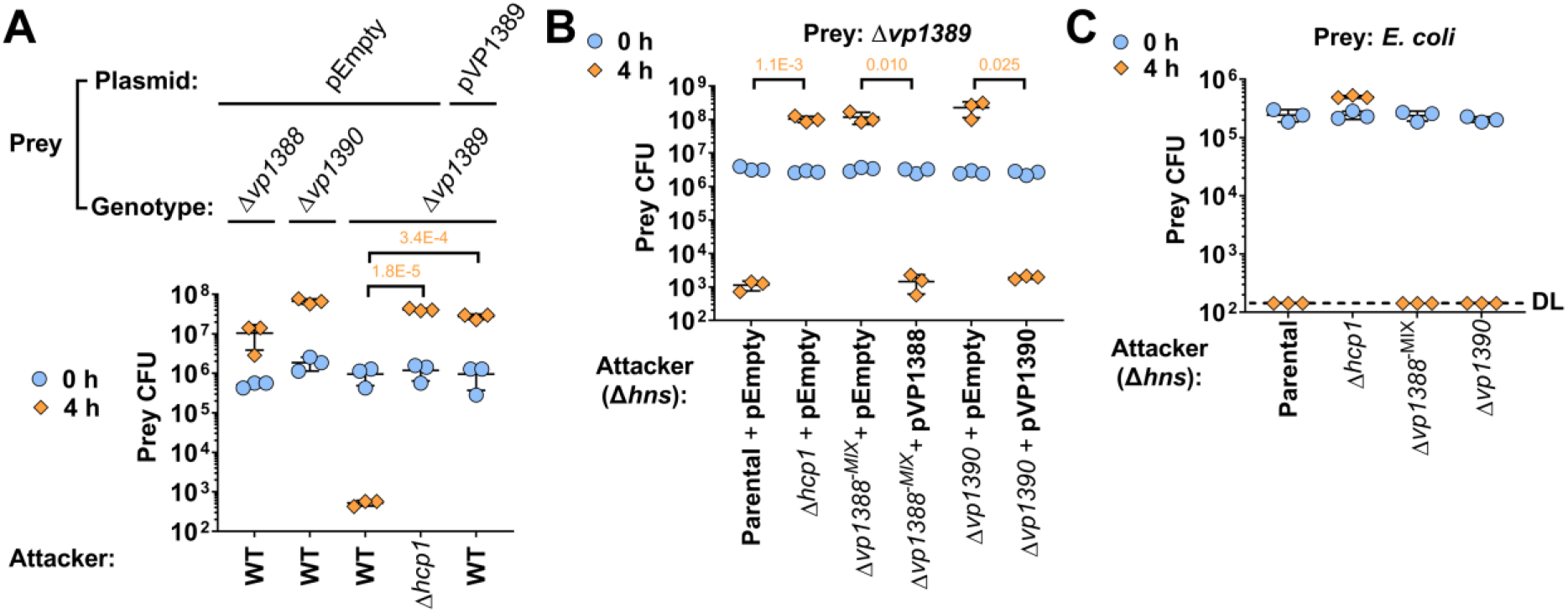
VP1388 and VP1390 are required for antibacterial toxicity, not immunity. **A-C)** Viability counts of the indicated *V. parahaemolyticus* (A-B) or *E. coli* (C) prey strains before (0 h) and after (4 h) co-incubation with the indicated *V. parahaemolyticus* attackers on media containing 3% NaCl at 30 °C. In A, prey strains contain either an empty plasmid (pEmpty) or a plasmid for arabinose-inducible expression of VP1389 (pVP1389). In B and C, prey strains contain an empty plasmid that provides a selection marker, and the attackers are derivatives of a *Δhns* mutant (parental). In B, the attackers contain an empty plasmid, or plasmids for the arabinose-inducible expression of VP1388 (pVP1388) or VP1390 (pVP1390). Data are shown as the mean ± SD. Statistical significance between samples at the 4 h timepoint by an unpaired, two-tailed Student’s *t*-test is denoted above. A significant difference was considered as *P* < 0.05. DL, assay detection limit. Δ*hcp1* was used as a T6SS1-control strain.

### Generating a *vp1388*^−^ mutant that does not affect VP1390 expression

Before performing additional experiments to investigate the tricistronic operon, we determined whether the single gene deletions that we used in Fig. 2A affected the expression of either VP1388 or VP1390. Although deletion of *vp1390* did not affect the expression of VP1388, deletion of *vp1388* resulted in elevated expression of VP1390 (Supplementary Fig. S2A). Since we did not wish to conduct subsequent experiments with a mutant in which the VP1390 expression levels are drastically elevated, we generated an alternative *vp1388* mutant in which the region encoding the MIX domain (corresponding to amino acids 242-423 (Salomon *et al*, 2014a)) was deleted. The resulting mutant, hereafter termed Δ*vp1388^−MIX^*, exhibited no detectable expression of VP1388 but retained VP1390 levels comparable to those of the wild-type strain (Supplementary Fig. S2A). Neither Δ*vp1388^−MIX^* nor the other single-gene deletion mutants revealed any growth defects (Supplementary Fig. S2B). Therefore, Δ*vp1388^−MIX^* was chosen to serve as a *vp1388^−^* strain in subsequent experiments. Surprisingly, VP1390 expression was absent in the Δ*vp1389* mutant (Supplementary Fig. S2A). We reasoned that this deletion resulted in a polar effect.

### VP1388 and VP1390 are both required for operon-mediated toxicity

We previously showed that VP1388 is required for T6SS1-mediated intoxication of a *vp1388-vp1389* deletion prey (Salomon *et al*, 2014a). Since we found no evidence of VP1390 playing a role in immunity, we hypothesized that it plays a role in the toxic activity mediated by the tricistronic operon. Indeed, competition assays revealed that both *vp1388^−^* (Δ*vp1388^−MIX^*) and *vp1390^−^* (Δ*vp1390*) mutants were unable to intoxicate the sensitive Δ*vp1389* prey, whereas exogenous expression of either VP1388 or VP1390 from a plasmid complemented the mutation (Fig. 2B). Notably, the attacker strains that were used for these assays were generated in a background in which *hns*, encoding a negative regulator of T6SS1 (Salomon *et al*, 2014b; Fridman *et al*, 2020), was deleted (Δ*hns*) to ensure maximal activation of T6SS1. The growth of Δ*hns* derivatives was comparable to that of their parental strain (Supplementary Fig. S3). Importantly, neither the *vp1388^−^* mutant nor the *vp1390^−^* mutant was impaired in its ability to intoxicate an *E. coli* prey, which unlike a Δ*vp1389* prey, is expected to be sensitive to toxicity mediated by other T6SS1 effector and immunity modules (Fig. 2C). These results indicate that VP1388 and VP1390 are both required for the toxic activity mediated by the tricistronic operon, but not for overall T6SS1 activity.

### VP1388 and VP1390 interact and are loaded onto the T6SS together

Since both VP1388 and VP1390 are secreted by T6SS1 (Salomon *et al*, 2014a) and are required for T6SS1-mediated toxicity (Fig. 2B), and since they are genetically linked (Fig. 1), we hypothesized that the two proteins physically interact. Indeed, immunoprecipitation assays of proteins co-expressed in *E. coli* confirmed that VP1390 specifically binds VP1388, whereas neither VP1390 nor VP1388 interacted with a control protein (Fig. 3A).

**Figure 3.**
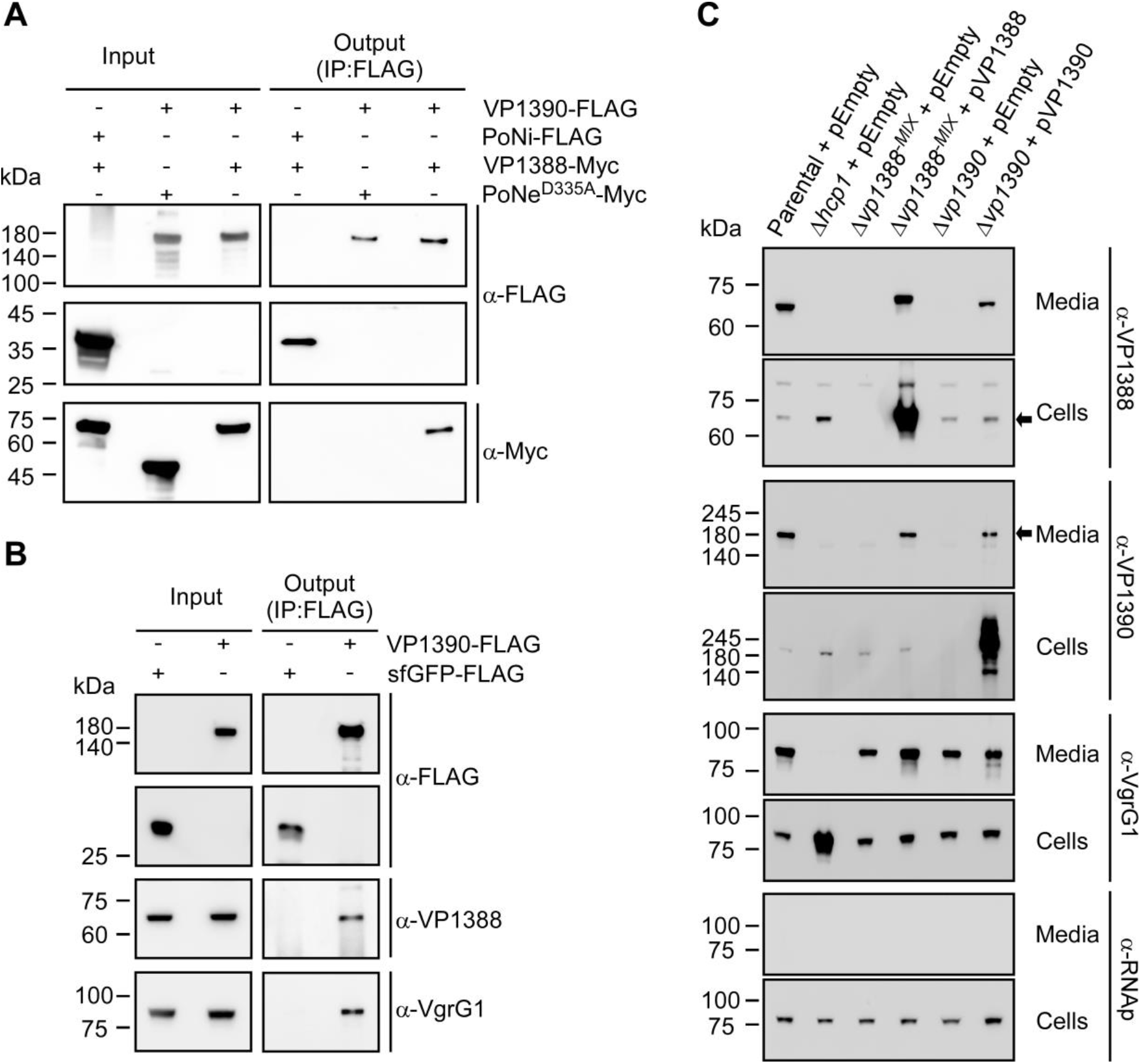
VP1388 and VP1390 are loaded together on the T6SS spike and are secreted co-dependently. **A)** VP1388 binds VP1388. Immunoprecipitation using α-FLAG antibodies from *E. coli* cells co-expressing the indicated C-terminal FLAG- and Myc-tagged proteins from arabinose-inducible plasmids. **B)** VP1388 and VgrG1 co-precipitate with VP1390. Immunoprecipitation using α-FLAG antibodies from *V. parahaemolyticus* Δ*hns*/Δ*hcp1*/Δ*vp1390* derivatives harboring plasmids for the arabinose-inducible expression of FLAG-tagged sfGFP or VP1390. Cells were grown in MLB media supplemented with chloramphenicol to maintain the plasmids, and 0.1% arabinose. Endogenous VP1388 and VgrG1 were detected using α-VP1388 and α-VgrG1 antibodies, respectively. **C)** Expression (cells) and secretion (media) of VP1388, VP1390, and VgrG1 from the indicated *V. parahaemolyticus* Δ*hns*-derived strains harboring an empty plasmid (pEmpty) or plasmids for the arabinose-inducible expression of VP1388 (pVP1388) or VP1390 (pVP1390). Samples were grown in media containing 3% NaCl and supplemented with 0.1% arabinose at 30 °C. RNA polymerase β (RNAp) was used as a non-secreted protein loading control.

For T6SS-mediated delivery, VP1388 and VP1390 must be loaded onto the T6SS tail tube. Considering their size, we reasoned that these proteins are not loaded into the narrow Hcp tube (Silverman *et al*, 2013), but rather, onto the spike comprising the VgrG and PAAR proteins (Nazarov *et al*, 2017). Therefore, we set out to determine whether VP1388 and VP1390 bind the T6SS1 spike in *V. parahaemolyticus*. To this end, we employed a strain in which *hcp1* was deleted; this was intended to prevent T6SS1-mediated secretion, which may result in losing a protein signal, while presumably retaining the assembly of the T6SS baseplate and spike (Brunet *et al*, 2015). As shown in Fig. 3B, immunoprecipitated VP1390, but not sfGFP that was used as a control, interacted with both VP1388 and VgrG1. This result suggests that VP1388 and VP1390 are loaded on the T6SS spike together.

### VP1388 and VP1390 are secreted co-dependently

We revealed that VP1388 and VP1390 interact, which led us to investigate whether their secretion is co-dependent. To this end, we monitored the secretion of VP1388 in the absence of VP1390 and *vice versa*. As shown in Fig. 3C, the secretion of VP1388 was abolished in the Δ*vp1390* strain and the secretion of VP1390 was abolished in the Δ*vp1388^−MIX^* strain; however, their expression was still detected in the absence of their counterpart, suggesting that they are not obligatory for each other’s expression and stability. Exogenous complementation of VP1388 or VP1390 from a plasmid restored their counterpart’s secretion. Notably, the absence of VP1388 or VP1390 did not affect the overall activity of T6SS1, since the secretion of the hallmark secreted spike protein, VgrG1, was retained in the Δ*vp1388^−MIX^* and Δ*vp1390* strains (Fig. 3C). Taken together, these results indicate that VP1388 and VP1390 form a heterocomplex that is required for their respective secretion via T6SS.

### VP1390 is an antibacterial toxin

Next, we set out to characterize the antibacterial activity of this operon and to determine which of the two secreted proteins mediates it. To this end, we investigated whether VP1388, VP1390, or both mediate antibacterial toxicity. Since the immunity protein, VP1389, contains an N-terminal signal peptide for periplasmic localization (Fig. 1A), we reasoned that the toxin will target this compartment. Therefore, we expressed VP1388 and VP1390, fused to an N-terminal PelB signal peptide (for periplasmic localization), in the surrogate host *E. coli* and monitored their effect on bacterial growth. Surprisingly, VP1390, but not VP1388, was toxic to *E. coli* (Fig. 4A). Expression of both VP1388 and VP1390 was detected by immunoblotting (Supplementary Fig. S4). This result suggests that VP1390 is the toxin responsible for the operon-mediated toxicity. In support of this notion, VP1390 specifically interacted with the immunity protein, VP1389, when both were exogenously co-expressed in *V. parahaemolyticus*, as expected from an effector and immunity pair (Fig. 4B). Notably, since over-expression of VP1389 itself was toxic in *E. coli*, we were unable to directly examine its ability to antagonize the toxicity mediated by VP1390 in this host.

**Figure 4.**
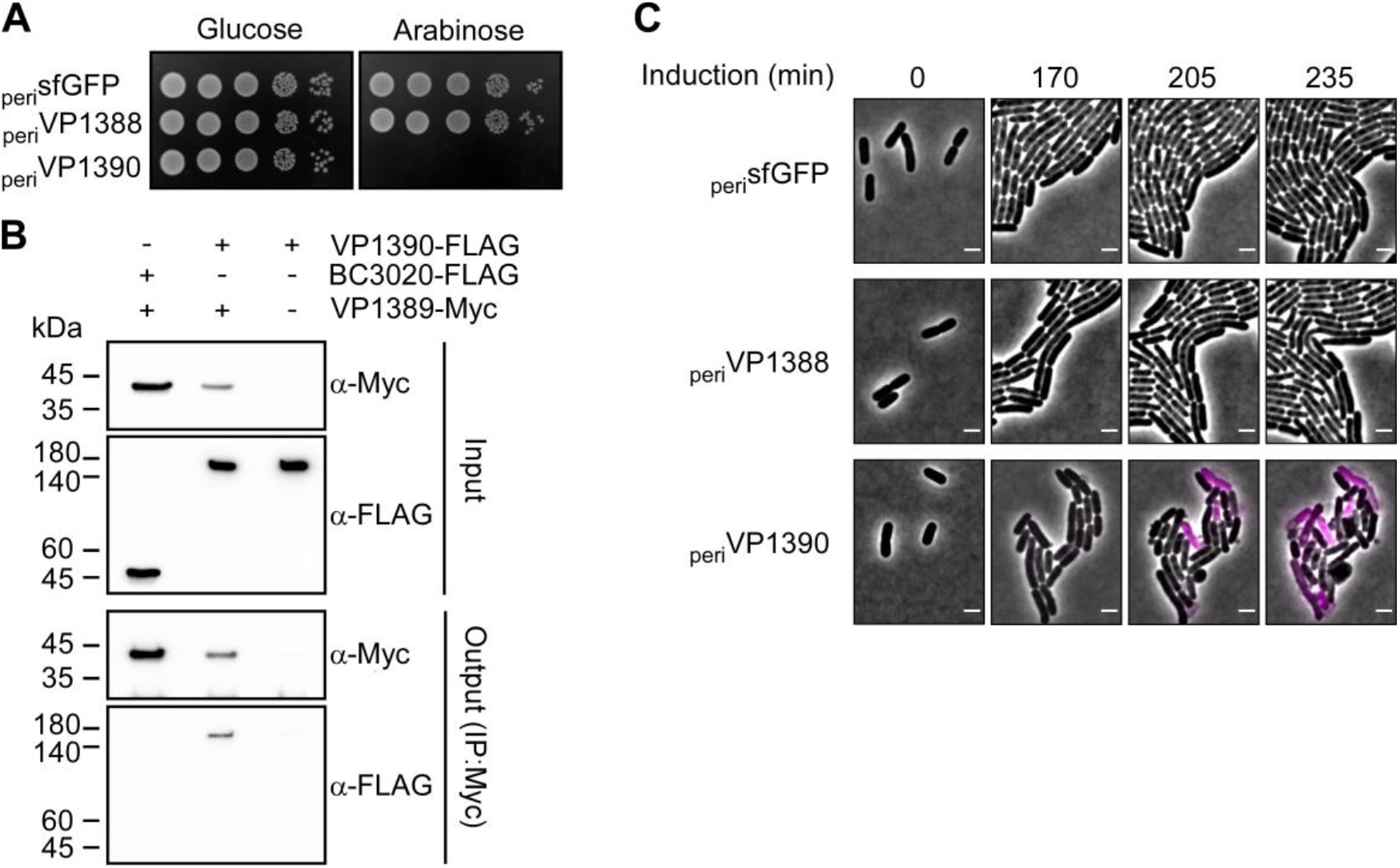
VP1390 is a periplasm-targeting toxin that leads to cell lysis. **A)** Toxicity of periplasm-targeted proteins in *E. coli*. *E. coli* strains containing plasmids for the arabinose-inducible expression of sfGFP (used as a control), VP1388 or VP1390 fused to an N-terminal PelB signal peptide (_peri_sfGFP, _peri_VP1388, and _peri_VP1390, respectively) were spotted at 10-fold serial dilutions onto LB agar plates supplemented with kanamycin (to maintain plasmids) and either 0.2% glucose, to repress protein expression, or 0.1% arabinose, to induce protein expression. **B)** VP1389 interacts with VP1390. Co-immunoprecipitation of FLAG-tagged VP1390 or BC3020 using Myc-tagged VP1389 when co-expressed in *V. parahaemolyticus* Δ*vp1389* (input). Precipitated proteins (output) were detected by immunobotting using α-Myc and α-FLAG antibodies. **C)** VP1390 induces cell lysis in *E. coli*. Time-lapse microscopy of *E. coli* cells expressing periplasm-targeted sfGFP, VP1388, or VP1390 (_peri_sfGFP, _peri_VP1388, and _peri_VP1390, respectively) from an arabinose-inducible vector, grown on LB agarose pads supplemented with kanamycin (to maintain the plasmid), 0.2% arabinose (to induce expression), and propidium iodide (PI; pink). Merging of the phase contrast and PI channels are shown. Scale bar = 2 μm.

To investigate the nature of the toxic activity mediated by VP1390, we monitored the morphological changes that occur in *E. coli* expressing VP1390. As shown in Fig. 4C and Supplementary Movie. S1, *E. coli* expressing periplasm-targeted VP1390, but not VP1388 or sfGFP, lysed and exhibited massive blebbing. Lysis was determined by changes in cell appearance as observed in the phase contrast channel, and by entry of the membrane-impermeable fluorescent DNA dye, propidium iodide.

### Triad induces T6SS1-mediated cell lysis in septating prey cells

To determine whether the lysis observed in *E. coli* expressing VP1390 is also mediated by the tricistronic operon during T6SS1-mediated competition, we monitored GFP-expressing, sensitive *V. parahaemolyticus* Δ*vp1389* prey cells during incubation with a T6SS1^+^ (Δ*hns*) or a T6SS1^−^ (Δ*hns*/Δ*hcp1*) attacker. As shown in Fig. 5 and Supplementary Movie. S2, Δ*vp1389* prey cells expressing GFP often lysed after contacting a T6SS1^+^ attacker cell. Often (62.17 ± 9.59%), lysis occurred in cells nearing completion of septation. Furthermore, when lysing cells were not crowded, a bleb containing cytoplasmic content (as indicated by the presence of GFP in it) often emerged from the septum prior to lysis and the entry of propidium iodide (Supplementary Fig. S5). Similar phenotypes were not observed in prey cells that were co-incubated with a T6SS1^−^ attacker, indicating that the lysis resulted from the T6SS1 triad activity. Taken together, these results indicate that VP1390 is the toxin component of the *vp1388*-*vp1390* triad, and that it leads to cell lysis upon delivery to the prey periplasm.

**Figure 5.**
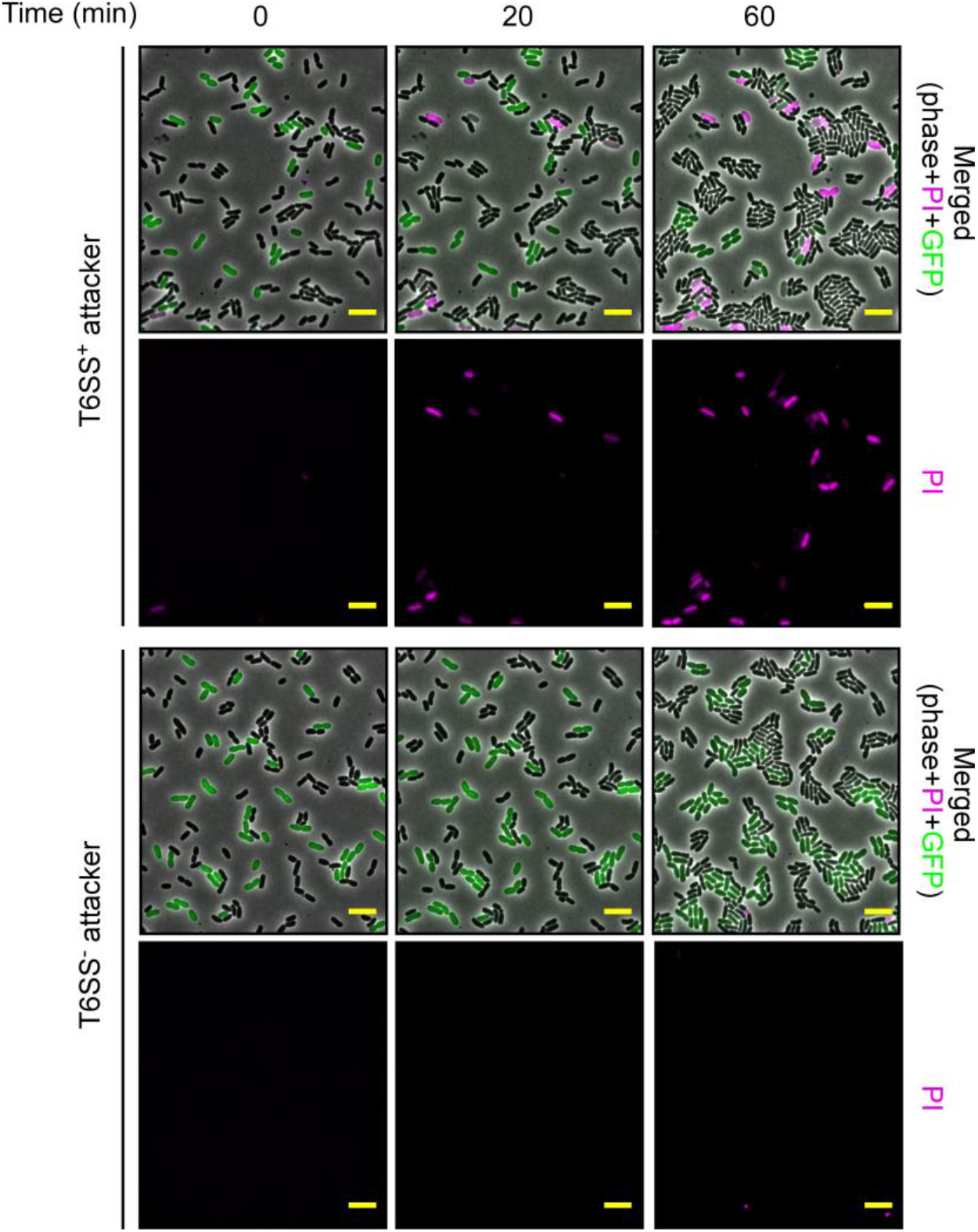
Operon-mediated toxicity results in prey cell lysis. Time-lapse microscopy of competition between *V. parahaemolyticus* Δ*hns* (T6SS+) or Δ*hns*/Δ*hcp1* (T6SS−) attackers and *V. parahaemolyticus* Δ*vp1389* prey that express GFP. Attacker and prey were mixed (2:1 ratio) and spotted on LB agarose pads supplemented with propidium iodide (PI; pink). Merging of the phase contrast, GFP (green), and PI (pink) channels, as well as the PI channel alone are shown. Scale bar = 5 μm.

## DISCUSSION

In this work, we characterized the role of the three proteins encoded in the *V. parahaemolyticus* T6SS-associated operon, *vp1388-90*, which was previously shown to mediate T6SS-dependent bacterial competition (Salomon *et al*, 2014a). Our results revealed a new mechanism underlying T6SS secretion in which VP1388, a MIX domain-containing protein, serves as a co-effector enabling the T6SS-mediated co-secretion of a novel antibacterial toxin, VP1390. We showed that VP1388 and VP1390 interact with each and are loaded on the T6SS spike; we also showed that they depend on each other for T6SS-mediated secretion. Therefore, we propose that VP1388 and VP1390 exemplify a previously undescribed mechanism of T6SS secretion, which we termed “a binary effector module”.

In previous works, we and others described diverse examples of polymorphic antibacterial and anti-eukaryotic T6SS toxins that contain a MIX domain (Bernal *et al*, 2017; Dar *et al*, 2018; Salomon *et al*, 2014a; Ray *et al*, 2017; Salomon *et al*, 2015; Miyata *et al*, 2011). The MIX-containing VP1388, however, does not appear to exert antibacterial toxicity as would be expected if it was the toxin responsible for the T6SS-dependent antibacterial toxicity mediated by T6SS1. Since VP1388 is required for secretion of VP1390, which does mediate antibacterial toxicity, we concluded that VP1388 plays another role for MIX domain-containing proteins as co-effectors, enabling the loading and secretion of toxins via T6SS. We hypothesize that VP1388 serves as a tether that connects the toxin, VP1390, to the T6SS spike, possibly to VgrG (Fig. 6). Nevertheless, although we have made numerous attempts to decipher the molecular mechanism that is used by VP1388 to enable VP1390 secretion, inconclusive results, possibly due to the “sticky” nature of the *V. parahaemolyticus* T6SS1 spike proteins, which we have experienced in some expression systems, prevent us from shedding more light on the mechanism in detail at this stage.

**Figure 6.**
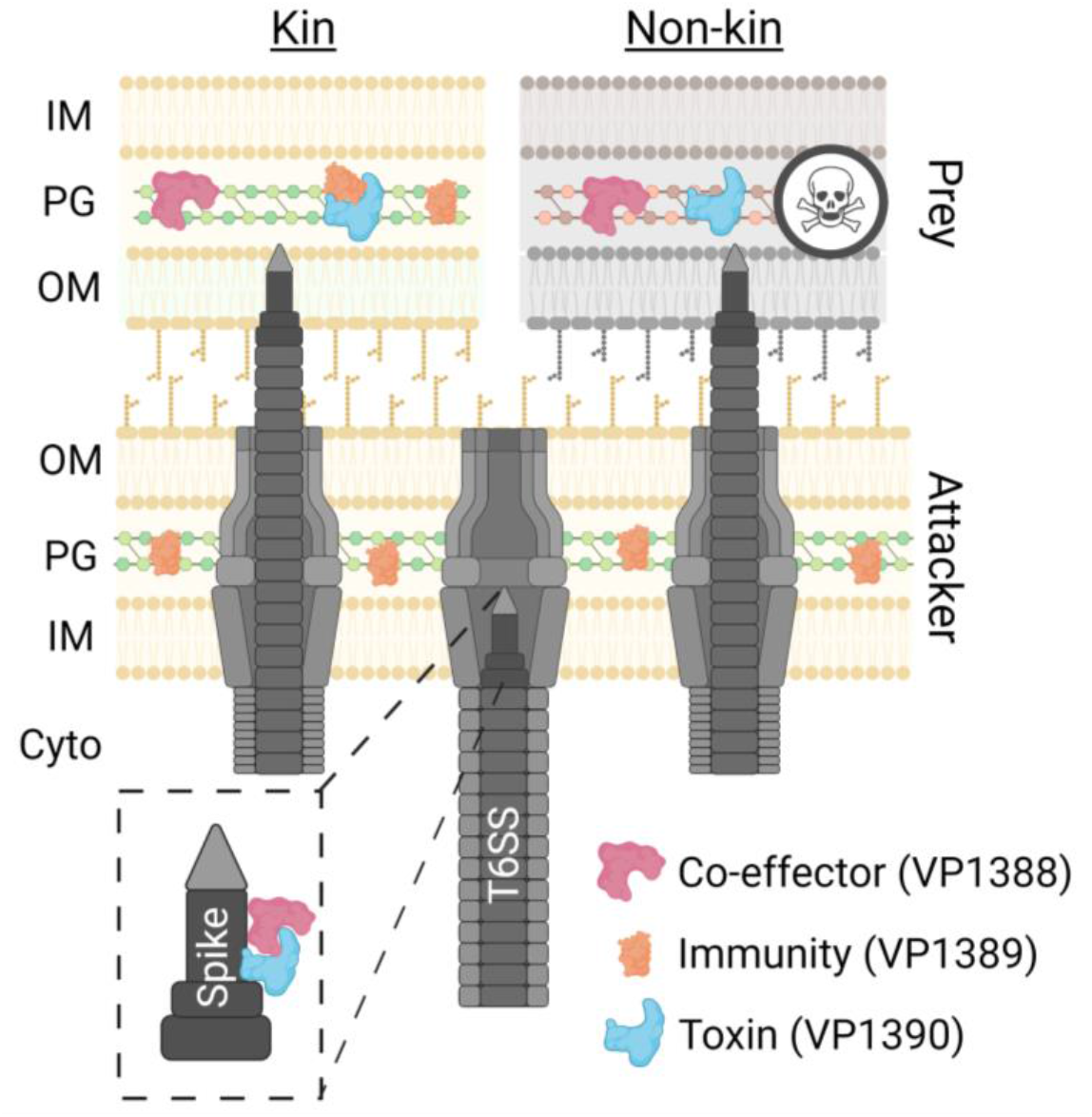
Model of T6SS binary effector delivery. The toxin, VP1390, and its co-effector, VP1388, are loaded together onto the T6SS spike and are delivered into the periplasm of a neighboring prey cell. If the prey cell expresses the cognate immunity protein, VP1389, then it can antagonize the attack (kin). Otherwise, the VP1390 toxin acts in the prey periplasm, leading to cell lysis (non-kin). IM, inner membrane; PG, peptidoglycan; OM, outer membrane; Cyto, cytoplasm. The figure was created using BioRender.com.

Proteins known as adaptors or chaperones were shown to interact with cognate effectors and T6SS tail tube components to stabilize and mediate the loading of effectors onto the T6SS (Manera *et al*, 2021). Nevertheless, we contend that VP1388 is not an adaptor or chaperone per se. First, VP1388 is secreted in a T6SS-dependent manner, whereas adaptors are not. An exception to this is the secreted tail tube component Hcp, which acts as a chaperone that stabilizes and delivers certain effectors (Silverman *et al*, 2013). However, Hcp, unlike VP1388, is a conserved and essential T6SS structural component. Second, VP1390 was stably expressed in *V. parahaemolyicus* even in the absence of VP1388, and it was toxic when expressed alone in *E. coli*. We cannot, however, rule out the possibility that VP1388 stabilizes at least some structural part of VP1390, which enables its proper loading onto the T6SS spike. Intriguingly, the absence of a VP1388 homolog in some bacterial species that encode VP1389 and VP1390-homologous dyads suggests that in these bacteria a different mechanism may be used to load the VP1390 homologs onto the T6SS spike for secretion.

VP1390 is a previously unrecognized antibacterial T6SS effector. It bears no resemblance to previously described toxins, aside from the OmpA_C-terminal-like domain, which is predicted to bind peptidoglycan (Koebnik, 1995). Indeed, we showed that VP1390 exerts its toxicity in the bacterial periplasm, leading to cell lysis. The morphological phenotypes observed during T6SS-mediated competition against *V. parahaemolyticus* prey lacking the periplasmic targeted immunity protein, VP1389, in which the cytosolic content was excreted in a bleb that often originated from the septum of cells nearing completion of division, suggest that VP1390 targets the peptidoglycan integrity. This is also supported by the lysis phenotype observed when VP1390 is expressed in the periplasm of *E. coli*. Future work will determine whether VP1390 indeed targets the peptidoglycan, and if so, whether it directly modifies the cell wall or whether it does so indirectly by manipulating proteins that regulate the cell wall.

The widespread nature of homologous tricistronic operons in marine bacteria, and their association with T6SSs emphasize their importance to the competitive fitness of these bacteria. Since many of these marine bacteria are established and emerging pathogens, better understanding the role of these genes will contribute to our ability to combat them. It remains to be determined whether other MIX domain-containing proteins serve as co-effectors in binary effector modules rather than as toxins per se.

In conclusion, in this work we revealed a previously undescribed T6SS effector secretion mechanism, whereby a co-effector that contains a MIX domain, previously thought to only be present in polymorphic toxins, enables the delivery of a toxin. We also characterized VP1390, a novel antibacterial toxin that induces bacterial cell lysis by a yet to be determined mechanism.

## MATERIALS AND METHODS

### Strains and media

For a complete list of strains used in this study, see Supplementary Table S1. *Escherichia coli* strains were grown in 2xYT broth (1.6% [wt/vol] tryptone, 1% [wt/vol] yeast extract, and 0.5% [wt/vol] NaCl) or Lysogeny broth (LB) at 37°C. Media were supplemented with kanamycin (30 μg/mL) or chloramphenicol (10 μg/mL) when appropriate to maintain plasmids. *Vibrio parahaemolyticus* was grown in MLB broth (LB containing 3% [wt/vol] NaCl) or on marine minimal media (MMM) agar plates (1.5% [wt/vol] agar, 2% [wt/vol] NaCl, 0.4% [wt/vol] galactose, 5 mM MgSO4, 7 mM K2SO4, 77 mM K2HPO4, 35 mM KH2PO4, and 2 mM NHCl) at 30°C. Media were supplemented with kanamycin (250 μg/mL) or chloramphenicol (10 μg/mL) when appropriate to maintain plasmids.

### Plasmid construction

For a complete list of plasmids used in this study, see Supplementary Table S2. Primers used for amplification are listed in Supplementary Table S3. For protein expression, the coding sequences (CDS) of the operon genes encoding NP_797767.1 (VP1388), NP_797768.1 (VP1389), and NP_797769.1 (VP1390) were amplified from *V. parahaemolyticus* strain RIMD 2210633 genomic DNA. The CDS of superfolder GFP (sfGFP) was amplified from the plasmid sfGFP-N1. Amplicons were inserted into the multiple cloning site (MCS) of pBAD/Myc-His, pBAD33.1 or their derivatives using the Gibson assembly method (Gibson *et al*, 2009) or by restriction digestion and ligation.

Plasmids were introduced into *E. coli* using electroporation or the Zymo Research MIX & Go kit, according to the manufacturer’s protocol. Transformants introduced with arabinose-inducible vectors were grown on agar plates supplemented with 0.2% [wt/vol] glucose to repress unwanted expression from the P*bad* promotor during the subcloning steps. Plasmids were introduced into *V. parahaemolyticus* via conjugation. Transconjugants were grown on MMM agar plates supplemented with appropriate antibiotics to maintain the plasmids.

### Construction of deletion strains

For in-frame deletions of the *vp1388* region encoding the MIX domain, *vp1389*, and *vp1390* from *V. parahaemolyticus* RIMD 2210633 genome, 1 kb upstream and 1 kb downstream of each gene or region to be deleted were amplified and cloned into pDM4, a Cm^R^Ori6k suicide plasmid (O’Toole *et al*, 1996) using restriction digestion and ligation. These vectors were transformed into electrocompetent *E. coli* S17-1 (λ pir) or DH5α (λ pir), and transferred into *V. parahaemolyticus* via conjugation. Transconjugants were first selected on MMM agar plates supplemented with chloramphenicol, and then transferred to MMM agar plates supplemented with sucrose (15% [wt/vol]) for counter-selection and loss of the SacB-containing pDM4. Deletions were confirmed by PCR. Construction of pDM4 plasmids for deletion of *vp1388* and *hns* (*vp1133*) was described previously (Salomon *et al*, 2014a, 2014b).

### Bacterial competition assays

Attacker and prey bacterial strains were grown overnight in MLB (*V. parahaemolyticus*) or LB (*E. coli*) broth supplemented with antibiotics when plasmid maintenance was required. Bacterial cultures were then normalized to OD_600_ = 0.5, and mixed at a 4:1 (attacker:prey) ratio. The mixtures were spotted on agar assay plates (MLB supplemented with 0.1% [wt/vol] L-arabinose to induce expression from plasmids) in triplicates and incubated at 30°C for 4 hours. Colony-forming units (CFU) of prey spotted at t = 0 hours were determined by plating 10-fold serial dilutions on selective agar plates. After 4 hours (t = 4 h), bacterial spots were scraped from assay agar plates into 1 mL of LB media. Next, 10-fold serial dilutions were spotted as described for t = 0 hours, and prey CFU were calculated. Assays were repeated three times with similar results; the results from a representative experiment are shown.

### Endogenous expression of VP1388 and VP1390 in *V. parahaemolyticus*

*V. parahaemolyticus* strains were grown overnight in MLB broth at 30°C. Overnight cultures were normalized to OD_600_ = 0.18 in 5 mL MLB supplemented with 20 μM phenamil (an inhibitor of the polar flagella used to mimic surface sensing activation) to induce the expression of the T6SS1 genes (Salomon *et al*, 2013). After 5 hours, 1.0 OD_600_ units of cells were pelleted and resuspended in (2X) Tris-Glycine SDS sample buffer (Novex, Life Sciences). Samples were boiled, and cell lysates were resolved on Mini-PROTEAN TGX Stain-Free™ precast gels (Bio-Rad) and transferred onto 0.2 μm nitrocellulose membranes. For immunoblotting, primary antibodies specific for VP1388 or VP1390 (α-VP1388 polyclonal antibody raised in rabbit against peptide CLAEDLQPVDKETQM, and α-VP1390 polyclonal antibody raised in rabbit against peptide EDENNDKTYPSWHSC, respectively; GenScript) were used at 1:1000 concentration. Protein signals were visualized in a Fusion FX6 imaging system (Vilber Lourmat) using enhanced chemiluminescence (ECL) reagents.

### *Vibrio* growth assays

Overnight cultures of *V. parahaemolyticus* strains were normalized to OD_600_ = 0.01 in MLB broth and transferred to 96-well plates (200 μL per well; n=4). The 96-well plates were incubated in a microplate reader (BioTek SYNERGY H1) at 30°C with constant shaking at 205 cpm. OD_600_ reads were acquired every 10 minutes.

### Toxicity in *E. coli*

*E. coli* strains carrying the indicated arabinose-inducible expression plasmids were grown in 2xYT broth supplemented with the appropriate antibiotics and 0.2% (wt/vol) glucose at 37°C. Overnight cultures were washed twice with fresh 2xYT broth to remove residual glucose. Cultures were then normalized to OD_600_ = 1 in 2xYT media supplemented with antibiotics. Next, 10-fold serial dilutions (dilutions 10^−1^-10^−5^) were spotted (5 μL) onto LB agar plates supplemented with antibiotics (to maintain plasmids) and 0.2% (wt/vol) glucose (to repress expression from the P*bad* promoter) or 0.1% (wt/vol) L-arabinose (to induce protein expression). Plates were incubated overnight at 30°C. The following morning, plates were imaged using a Fusion FX6 imaging system (Vilber Lourmat).

### Protein expression in *E. coli*

*E. coli* strains containing arabinose-inducible plasmids for C-terminal Myc-tagged protein expression were grown in 2xYT broth supplemented with appropriate antibiotics and 0.2% (wt/vol) glucose at 37°C. Overnight cultures were washed twice with fresh 2xYT broth to remove residual glucose. Cultures were then normalized to OD_600_ = 0.5 in 3 mL 2xYT broth supplemented with appropriate antibiotics and grown for two hours at 37°C. After 2 hours, 0.1% (wt/vol) L-arabinose was added to the media to induce protein expression, and cultures were grown for 2 additional hours at 37°C. Following induction, 0.5 OD_600_ units of cells were pelleted and resuspended in (2X) Tris-Glycine SDS sample buffer (Novex, Life Sciences). Samples were boiled, and cell lysates were resolved on TGX Stain-Free™ precast gels (Bio-Rad) and analyzed as mentioned above. For immunoblotting, α-Myc antibodies (Santa Cruz Biotechnology, 9E10, mouse mAb) were used at 1:1000 dilution.

### *E. coli* immunoprecipitation assays

To identify the direct interaction between VP1388 and VP1390, *E. coli* BL21 (DE3) cells harboring pBAD33.1-based plasmids encoding C-terminal FLAG-tagged PoNi (B5C30_RS14460) (Jana *et al*, 2019) or VP1390, together with pBAD/Myc-His-based plasmids encoding VP1388 or PoNe^D335A^ (B5C30_RS14465) (Jana *et al*, 2019) were grown overnight in 2xYT media supplemented with kanamycin and chloramphenicol at 37°C. Overnight cultures were diluted 1:100 in 50 mL fresh 2xYT media supplemented with appropriate antibiotics, and grown at 37°C for 2 h. After 2 h, 0.1% (wt/vol) L-arabinose was added to induce protein expression, and cultures were further grown at 30°C for 4 h. Next, 200 OD_600_ units were pelleted by centrifugation at 3,500 × *g* for 10 minutes at 4°C. Then, cell pellets were resuspended in 3 mL of Lysis buffer C (150 mM NaCl, 20 mM Tris-HCl pH = 7.5, 1 mM EDTA, and 0.5% [vol/vol] NP-40) supplemented with 0.1 mM PMSF, and were lysed using a high-pressure homogenizer (Multi cycle cell disruptor, Constant Systems). Cell debris was removed by centrifuging at 15,000 × *g* for 20 minutes at 4°C. Next, 500 μL of supernatant were mixed with 10 μL of DYKDDDDK Tag antibody (α-FLAG) and incubated for 1 hour at 4°C with constant rotation. Next, protein A and protein G magnetic beads (12.5 μL each) were mixed and prewashed with Lysis buffer C, and then mixed with the samples and incubated for an additional hour at 4°C with constant rotation. Beads were washed eight times with Lysis buffer C (200 μL each time). Finally, the beads were collected, and bound proteins were eluted by adding 100 μL of (2X) Tris-Glycine SDS Sample Buffer supplemented with 5% β-mercaptoethanol, followed by heating at 70°C for 5 minutes. Samples were analyzed by immunoblotting as mentioned above. HRP-conjugated α-Light Chain-specific secondary antibodies (Jackson ImmunoResearch) were used to avoid detecting the primary antibodies’ heavy chains.

### *Vibrio* immunoprecipitation assays

To detect loading of VP1390 and VP1388 on the T6SS spike, *V. parahaemolyticus* RIMD 2210633 Δ*hns*/Δ*hcp1*/Δ*vp1390* carrying the indicated pBAD33.1-based plasmids for expression of sfGFP or VP1390 with a C-terminal FLAG tag were grown overnight in MLB broth supplemented with chloramphenicol at 30°C. Overnight cultures were normalized to OD_600_ = 0.18 in 50 mL MLB broth supplemented with antibiotics and 0.1% (wt/vol) L-arabinose (to induce protein expression), and were grown at 30°C for 4 hours. After 4 hours, 140 OD_600_ units were pelleted at 3,500 × *g* for 10 minutes at 4°C. Then, 3.5 mL of Lysis buffer A (50 mM NaCl, 10 mM Tris-HCl pH = 7.5, 1 mM EDTA, 0.5% [vol/vol] NP-40, and 0.1 mM PMSF) were added to cell pellets, which were then incubated with rotation at 4°C for 15 minutes to resuspend the cells. Cells were then lysed using a high-pressure homogenizer (Multi cycle cell disruptor, Constant Systems). Cell debris was removed by centrifugation at 15,000 × *g* for 20 minutes at 4°C. Next, 490 μL of supernatant were incubated with 10 μL of DYKDDDDK Tag antibody (α-FLAG) for an hour at room temperature (RT). Protein A and protein G magnetic beads (25 μL and 10 μL, respectively), prewashed with Wash buffer A (50 mM NaCl, 10 mM Tris-HCl pH=7.5, 1 mM EDTA, and 0.5% [vol/vol] NP-40) were added to samples and incubated with constant rotation for an additional hour at RT. Then, samples were washed three times with Wash buffer, and the beads were collected. Bound proteins were eluted by adding 50 μL of (2X) Tris-Glycine SDS Sample Buffer supplemented with 5% β-mercaptoethanol, followed by heating at 70°C for 5 minutes. Samples were analyzed by immunoblotting as mentioned above. HRP-conjugated α-Light Chain-specific secondary antibodies (Jackson ImmunoReserach) were used to avoid detecting the primary antibodies’ heavy chains.

To detect the binding of VP1389 to VP1390, *V. parahaemolyticus* RIMD 2210633 Δ*vp1389* (which does not express endogenous VP1389 or VP1390, as shown in Supplementary Fig. S2A) carrying pBAD/Myc-His-based plasmids, either empty or encoding VP1389, together with pBAD33.1-based plasmids encoding C-terminally FLAG-tagged VP1390 or BC3020 (accession number NP_832766.1; used as control), were grown overnight in MLB broth supplemented with chloramphenicol and kanamycin at 30°C. Overnight cultures were normalized to OD_600_ = 0.18 in 50 mL MLB broth supplemented with antibiotics and 0.1% (wt/vol) L-arabinose (to induce protein expression), and were grown at 30°C for 3 hours. After 3 hours, 100 OD_600_ units were pelleted at 3,500 × *g* for 10 minutes at 4°C. Next, 3 mL of Lysis buffer B (100 mM NaCl, 10 mM Tris-HCl pH=7.5, 1 mM EDTA, 0.5% [vol/vol] NP-40, and 0.1 mM PMSF) were added to cell pellets, and cells were lysed, as detailed above. Cell debris was removed as mentioned above. Next, 500 μL of supernatant were transferred to tubes containing 25 μL of prewashed magnetic α-Myc beads (Myc-tag [9B11] mouse mAb magnetic beads conjugated #5698; Cell Signaling Technology) and incubated at 4°C for 2 hours. Then, samples were washed 3 times with Wash buffer B (100 mM NaCl, 10 mM Tris-HCl pH=7.5, 1 mM EDTA, and 0.5% [vol/vol] NP-40), and bound proteins were eluted by adding 50 μL of (2X) Tris-Glycine SDS Sample Buffer supplemented with 5% β-mercaptoethanol, followed by heating at 70°C for 5 minutes. Samples were analyzed by immunoblotting as mentioned above.

### Secretion assays

*V. parahaemolyticus* strains were grown overnight in MLB broth supplemented with antibiotics to maintain plasmids, when needed. Cultures were normalized to OD_600_ = 0.18 in 5 mL MLB supplemented with antibiotics and L-arabinose (0.1% [wt/vol]) to induce expression from P*bad* promoters. After 5 hours, 1.0 OD_600_ units were collected for expression fractions (cells). The cell pellets were resuspended in (2X) Tris-Glycine SDS sample buffer (Novex, Life Sciences). For secretion fractions (media), 10 OD_600_ units were filtered (0.22 μm), and proteins were precipitated from the media using deoxycholate and trichloroacetic acid (Bensadoun & Weinstein, 1976). Cold acetone was used to wash the protein precipitates twice. Then, protein precipitates were resuspended in 20 μL of 10 mM Tris-HCl pH=8, followed by the addition of 20 μL of (2X) Tris-Glycine SDS Sample Buffer supplemented with 5% β-mercaptoethanol. Next, 0.5 μL of 1 N NaOH was added to maintain a basic pH. Expression and secretion samples were boiled and then resolved on Mini-PROTEAN or Criterion™TGX Stain-Free™ precast gels (Bio-Rad) and analyzed as mentioned above. For immunoblotting, primary antibodies were used at 1:1000 concentration. The following antibodies were used: DYKDDDDK Tag Antibody (D6W5B rabbit mAb #14793; Cell Signaling Technology; it binds to the same epitope as Sigma’s Anti-FLAG M2 Antibody; it is referred to as α-FLAG), Direct-Blot™ HRP anti-*E. coli* RNA Sigma 70 (mouse mAb #663205; BioLegend; it is referred to as α-RNAp), custom-made α-VgrG1 (Li *et al*, 2017), α-VP1388 (described above), and α-VP1390 (described above). α-RNAp was used to determine equal loading of samples and to exclude cell lysis. Protein signals were visualized in a Fusion FX6 imaging system (Vilber Lourmat) using enhanced chemiluminescence (ECL) reagents.

### Microscopy

To determine the effect of protein expression in *E. coli*, overnight *E. coli* MG1655-derivative sAJM.1506 cells carrying pPER5-based plasmids were diluted 100-fold into 3 mL of fresh LB broth supplemented with kanamycin and 0.2% (wt/vol) glucose. After 2 hours of incubation at 37°C, cells were washed and normalized to OD_600_ = 0.5. Next, 1 μL of each culture was spotted on LB agarose pads (1% [wt/vol] agarose supplemented with 0.2% [wt/vol] L-arabinose) onto which 1 μL of the membrane-impermeable DNA dye, propidium iodide (PI; 1 mg/mL; Sigma) had been pre-applied. After the spots had dried (1-2 minutes at RT), the agarose pads were mounted, facing down, on 35 mm glass bottom CELLview™ cell culture dishes (Greiner). Cells were then imaged every 5 minutes for 4 hours under a fluorescence microscope, as detailed below. The stage chamber (Okolab) temperature was set to 37°C.

To assess the T6SS1-dependent toxic effect of the tricistronic operon on sensitive prey during bacterial competition, *V. parahaemolyticus* RIMD 2210633 Δ*vp1389* prey cells harboring a plasmid for the constitutive expression of GFP (Ritchie *et al*, 2012) were competed against *V. parahaemolyticus* RIMD 2210633 attacker strain Δ*hns* (T6SS1^+^) or Δ*hns*/Δ*hcp1* (T6SS1^−^). Bacteria were grown overnight in MLB broth at 30°C. Overnight attacker and prey cultures were diluted 100-fold into 3 mL of fresh MLB broth and grown for 2 hours at 30°C. After 2 hours, attacker and prey cultures were normalized to OD_600_ = 2.5 and mixed in a 2:1 (attacker:prey) ratio. Cell mixtures and PI were spotted onto agarose pads and processed as detailed above. Bacteria were imaged every 4 minutes for 1 hour. The stage chamber temperature was set to 30°C.

The following setup was used for imaging: a Nikon Eclipse Ti2E inverted motorized microscope with a CFI PLAN apochromat DM 100X oil lambda PH-3 (NA, 1.45) objective lens, a Lumencor SOLA SE II 395 light source, and ET-dsRED (#49005, CHROMA, to visualize the PI signal) and ET-EGFP (#49002, CHROMA, to visualize the GFP signal) filter sets and a DS-QI2 Mono cooled digital microscope camera (16 MO). The obtained images were further processed and analyzed using Fiji ImageJ suite (Schindelin *et al*, 2012).

### Construction of position-specific scoring matrices for VP1388, VP1389, and VP1390

The position-specific scoring matrices (PSSMs) of VP1388, VP1389, and VP1390 were constructed using full-length sequences from *Vibrio parahaemolyticus* RIMD 2210633 (BAC59651.1, BAC59652.1, and BAC59653.1, respectively). The PSSM of a distant homolog of VP1389 was constructed using the full-length sequence from *Vibrio parahaemolyticus* ISF-77-01 (WP_047482080.1). Five iterations of PSI-BLAST were performed against the RefSeq protein database. In each iteration, a maximum of 500 hits with an expect value threshold of 10^−6^ and a query coverage of 70% were used.

### Identification of homologous operons of *vp1388-90*

Homologous operons of *vp1388-vp1390* were identified by searching for homologs of VP1388 and VP1390 in bacterial genomes. A local database containing the RefSeq bacterial nucleotide and protein sequences was generated (last updated on December 25, 2020). RPS-BLAST was used to identify VP1388 and VP1390 homologs in the local database. The results were filtered using an expect value threshold of 10^−15^ and a subject coverage of 70%. Subsequently, the genomic neighborhood was analyzed as described before (Dar *et al*, 2018; Fridman *et al*, 2020). Duplicated protein accessions appearing in the same genome in more than one genomic accession were removed if the same downstream protein existed at the same distance. The obtained list represented all occurrences of the VP1388 and VP1390 homologs in bacterial genomes. A list of homologous operons was generated by collecting all occurrences of VP1388 homologs and the occurrences of VP1390 homologs that were not found within 5 genes downstream of VP1388.

### Identification of VP1388-VP1390 triads and dyads

The list of homologous operons was analyzed. For VP1388 homologs, the following rules were applied: (1) if a VP1390 homolog was identified 2 to 5 genes downstream of the VP1388 homolog, it was termed ‘triad’; (2) if a VP1389 homolog was identified 1 gene downstream of the VP1388 homolog, it was termed ‘other’; (3) otherwise, it was termed ‘truncated/pseudo’. For VP1390 homologs, the following rules were applied: (1) if a VP1388 homolog was identified 2 to 5 genes upstream of the VP1390 homolog, it was termed ‘triad’; (2) if a VP1389 homolog was identified 1 gene upstream of the VP1390 homolog and all of the 2 to 5 genes upstream existed and were unrelated to VP1388, it was termed ‘dyad’; (3) if a VP1388 was identified 1 gene upstream of the VP1390 homolog, it was termed ‘other’; (4) if all of the 5 genes upstream and downstream existed and were unrelated to VP1388 and VP1389, indicating that the VP1390 homolog was an orphan, it was termed ‘other’; (5) otherwise, it was termed ‘truncated/pseudo’. All annotations were assessed manually. Changes were noted in the appropriate Supplementary Dataset.

### Identification of bacterial genomes encoding T6SS

RPS-BLAST was employed to identify the T6SS core components, as described before (Jana *et al*, 2019). Briefly, the proteins were aligned against 11 COGs that were previously shown to specifically predict T6SS and were against COG3501 (VgrG) (Boyer *et al*, 2009). Bacterial genomes encoding at least 9 out of the 11 T6SS core components were identified.

BLASTX was employed to identify *Vibrio parahaemolyticus* T6SS1-like cluster proteins in bacterial genomes, as described before (Fridman *et al*, 2020). Briefly, translated nucleotide sequences were aligned against the 24 T6SS1 cluster proteins of *Vibrio parahaemolyticus* RIMD 2210633 (NP_797770.1 to NP_797793.1). The minimal similarity percentage (the bit-score value divided by two times the specific lengths of the cluster proteins) of each protein was defined as 50%. Bacterial genomes encoding at least 12 out of the 24 T6SS1 cluster proteins were regarded as harboring a *Vibrio parahaemolyticus* T6SS1-like cluster. Genomes containing less than 17 of the 24 genes were also evaluated manually.

### Construction of the phylogenetic tree of bacterial strains containing VP1388 and VP1390

Phylogenetic analysis was conducted using the MAFFT server (mafft.cbrc.jp/alignment/server/). DNA sequences of *rpoB* coding for DNA-directed RNA polymerase subunit beta were aligned using MAFFT v7 FFT-NS-2 (Katoh *et al*, 2018, 2002). Partial and pseudogene sequences were not included in the analysis. The evolutionary history was inferred using the neighbor-joining method (Saitou & Nei, 1987) with the Jukes-Cantor substitution model (JC69). The analysis included 1543 nucleotide sequences and 3912 conserved sites.

## ACKNOWLEDGMENTS

This project received funding from the European Research Council (ERC) under the European Union’s Horizon 2020 research and innovation program (Grant agreement No. 714224), and the Israel Science Foundation (ISF; grant no. 920/17) to DS. We thank members of the Salomon lab for technical assistance and helpful discussions.

## AUTHOR CONTRIBUTIONS

Conceptualization: YD, BJ, EB, and DS; Methodology: YD, BJ, EB, and DS; Investigation: YD, BJ, and EB; Supervision: DS; Writing—original draft: DS; Writing—review & editing: YD, BJ, EB, and DS.

## REFERENCES

Ahmad S, Tsang KK, Sachar K, Quentin D, Tashin TM, Bullen NP, Raunser S, McArthur AG, Prehna G & Whitney JC (2020) Structural basis for effector transmembrane domain recognition by type vi secretion system chaperones. Elife 9: 1–29

Alcoforado Diniz J & Coulthurst SJ (2015) Intraspecies Competition in Serratia marcescens Is Mediated by Type VI-Secreted Rhs Effectors and a Conserved Effector-Associated Accessory Protein. J Bacteriol 197: 2350–60

Basler M, Pilhofer M, Henderson GP, Jensen GJ & Mekalanos JJ (2012) Type VI secretion requires a dynamic contractile phage tail-like structure. Nature 483: 182–6

Bensadoun A & Weinstein D (1976) Assay of proteins in the presence of interfering materials. Anal Biochem 70: 241–250

Bernal P, Allsopp LP, Filloux A & Llamas MA (2017) The Pseudomonas putida T6SS is a plant warden against phytopathogens. ISME J 11: 972–987

Berni B, Soscia C, Djermoun S, Ize B & Bleves S (2019) A type VI secretion system trans-kingdom effector is required for the delivery of a novel antibacterial toxin in Pseudomonas aeruginosa. Front Microbiol 10: 1218

Bondage DD, Lin J-S, Ma L-S, Kuo C-H & Lai E-M (2016) VgrG C terminus confers the type VI effector transport specificity and is required for binding with PAAR and adaptor–effector complex. Proc Natl Acad Sci 113: E3931–E3940

Boyd EF, Carpenter MR, Chowdhury N, Cohen AL, Haines-Menges BL, Kalburge SS, Kingston JJ, Lubin JBB, Ongagna-Yhombi SY & Whitaker WB (2015) Post-genomic analysis of members of the family Vibrionaceae. 3: 1–26

Boyer F, Fichant G, Berthod J, Vandenbrouck Y & Attree I (2009) Dissecting the bacterial type VI secretion system by a genome wide in silico analysis: What can be learned from available microbial genomic resources? BMC Genomics 10

Brunet YR, Zoued A, Boyer F, Douzi B & Cascales E (2015) The type VI secretion TssEFGK-VgrG phage-like baseplate is recruited to the TssJLM membrane complex via multiple contacts and serves as assembly platform for tail tube/sheath polymerization. PLOS Genet 11: e1005545

Burkinshaw BJ, Liang X, Wong M, Le ANH, Lam L & Dong TG (2018) A type VI secretion system effector delivery mechanism dependent on PAAR and a chaperone-co-chaperone complex. Nat Microbiol 3: 632–640

Cianfanelli FR, Alcoforado Diniz J, Guo M, De Cesare V, Trost M & Coulthurst SJ (2016) VgrG and PAAR proteins define distinct versions of a functional type VI secretion system. PLoS Pathog 12: 1–27

Dar Y, Salomon D & Bosis E (2018) The antibacterial and anti-eukaryotic Type VI secretion system MIX-effector repertoire in Vibrionaceae. Mar Drugs 16: 433

Flaugnatti N, Le TTH, Canaan S, Aschtgen M-S, Nguyen VS, Blangy S, Kellenberger C, Roussel A, Cambillau C, Cascales E, et al (2016) A phospholipase A 1 antibacterial Type VI secretion effector interacts directly with the C-terminal domain of the VgrG spike protein for delivery. Mol Microbiol 99: 1099–1118

Flaugnatti N, Rapisarda C, Rey M, Beauvois SG, Nguyen VA, Canaan S, Durand E, Chamot-Rooke J, Cascales E, Fronzes R, et al (2020) Structural basis for loading and inhibition of a bacterial T6 <scp>SS</scp> phospholipase effector by the VgrG spike. EMBO J 39

Fridman CM, Keppel K, Gerlic M, Bosis E & Salomon D (2020) A comparative genomics methodology reveals a widespread family of membrane-disrupting T6SS effectors. Nat Commun 11: 1085

Gibson DG, Young L, Chuang RY, Venter JC, Hutchison CA & Smith HO (2009) Enzymatic assembly of DNA molecules up to several hundred kilobases. Nat Methods 6: 343–345

Hachani A, Allsopp LP, Oduko Y & Filloux A (2014) The VgrG proteins are ‘à la carte’ delivery systems for bacterial type VI effectors. J Biol Chem 289: 17872–84

Hood RD, Singh P, Hsu FS, Güvener T, Carl MA, Trinidad RRS, Silverman JM, Ohlson BB, Hicks KG, Plemel RL, et al (2010) A type VI secretion system of Pseudomonas aeruginosa targets a toxin to bacteria. Cell Host Microbe 7: 25–37

Horseman MA, Bray R, Lujan-Francis B & Matthew E (2013) Infections Caused by Vibrionaceae. Infect Dis Clin Pract 21: 222–232

Hubert CL & Michell SL (2020) A universal oyster infection model demonstrates that *Vibrio vulnificus* <scp>Type 6</scp> secretion systems have antibacterial activity *in vivo*. Environ Microbiol 22: 4381–4393

Jana B, Fridman CM, Bosis E & Salomon D (2019) A modular effector with a DNase domain and a marker for T6SS substrates. Nat Commun 10: 3595

Katoh K, Misawa K, Kuma K & Miyata T (2002) MAFFT: a novel method for rapid multiple sequence alignment based on fast Fourier transform. Nucleic Acids Res 30: 3059–66

Katoh K, Rozewicki J & Yamada KD (2018) MAFFT online service: Multiple sequence alignment, interactive sequence choice and visualization. Brief Bioinform 20: 1160–1166

Koebnik R (1995) Proposal for a peptidoglycan-associating alpha-helical motif in the C-terminal regions of some bacterial cell-surface proteins. Mol Microbiol 16: 1269–1270

Lai H-C, Ng TH, Ando M, Lee C-T, Chen I-T, Chuang J-C, Mavichak R, Chang S-H, Yeh M-D, Chiang Y-A, et al (2015) Pathogenesis of acute hepatopancreatic necrosis disease (AHPND) in shrimp. Fish Shellfish Immunol 47: 1006–1014

Li P, Kinch LN, Ray A, Dalia AB, Cong Q, Nunan LM, Camilli A, Grishin N V, Salomon D & Orth K (2017) Acute hepatopancreatic necrosis disease-causing Vibrio parahaemolyticus strains maintain an antibacterial type VI secretion system with versatile effector repertoires. Appl Environ Microbiol 83: e00737–17

Liang X, Moore R, Wilton M, Wong MJQ, Lam L & Dong TG (2015) Identification of divergent type VI secretion effectors using a conserved chaperone domain. Proc Natl Acad Sci 112: 9106–9111

Ma J, Pan Z, Huang J, Sun M, Lu C & Yao H (2017) The Hcp proteins fused with diverse extended-toxin domains represent a novel pattern of antibacterial effectors in type VI secretion systems. Virulence 8: 1189–1202

MacIntyre DL, Miyata ST, Kitaoka M & Pukatzki S (2010) The Vibrio cholerae type VI secretion system displays antimicrobial properties. Proc Natl Acad Sci 107: 19520–19524

Manera K, Kamal F, Burkinshaw B & Dong TG (2021) Essential functions of chaperones and adaptors of protein secretion systems in Gram-negative bacteria. FEBS J

Miyata ST, Kitaoka M, Brooks TM, McAuley SB & Pukatzki S (2011) Vibrio cholerae requires the type VI secretion system virulence factor vasx to kill dictyostelium discoideum. Infect Immun 79: 2941–2949

Mougous JD, Cuff ME, Raunser S, Shen A, Zhou M, Gifford CA, Goodman AL, Joachimiak G, Ordoñez CL, Lory S, et al (2006) A virulence locus of Pseudomonas aeruginosa encodes a protein secretion apparatus. Science (80-) 312: 1526–1530

Nazarov S, Schneider JP, Brackmann M, Goldie KN, Stahlberg H & Basler M (2017) Cryo‐EM reconstruction of Type VI secretion system baseplate and sheath distal end. EMBO J 37: e201797103

Newton A, Kendall M, Vugia DJ, Henao OL & Mahon BE (2012) Increasing Rates of Vibriosis in the United States, 1996–2010: Review of Surveillance Data From 2 Systems. Clin Infect Dis 54: S391–S395

O’Toole R, Milton DL & Wolf-Watz H (1996) Chemotactic motility is required for invasion of the host by the fish pathogen Vibrio anguillarum. Mol Microbiol 19: 625–637

Pukatzki S, Ma AT, Revel AT, Sturtevant D & Mekalanos JJ (2007) Type VI secretion system translocates a phage tail spike-like protein into target cells where it cross-links actin. Proc Natl Acad Sci 104: 15508–15513

Pukatzki S, Ma AT, Sturtevant D, Krastins B, Sarracino D, Nelson WC, Heidelberg JF & Mekalanos JJ (2006) Identification of a conserved bacterial protein secretion system in Vibrio cholerae using the Dictyostelium host model system. Proc Natl Acad Sci 103: 1528–1533

Quentin D, Ahmad S, Shanthamoorthy P, Mougous JD, Whitney JC & Raunser S (2018) Mechanism of loading and translocation of type VI secretion system effector Tse6. Nat Microbiol 3: 1142–1152

Ray A, Schwartz N, Souza Santos M, Zhang J, Orth K, Salomon D, de Souza Santos M, Zhang J, Orth K & Salomon D (2017) Type VI secretion system MIX‐effectors carry both antibacterial and anti‐eukaryotic activities. EMBO Rep 18: e201744226

Ritchie JM, Rui H, Zhou X, Iida T, Kodoma T, Ito S, Davis BM, Bronson RT & Waldor MK (2012) Inflammation and Disintegration of Intestinal Villi in an Experimental Model for Vibrio parahaemolyticus-Induced Diarrhea. PLoS Pathog 8: e1002593

Russell AB, Hood RD, Bui NK, Leroux M, Vollmer W & Mougous JD (2011) Type VI secretion delivers bacteriolytic effectors to target cells. Nature 475: 343–349

Russell AB, Singh P, Brittnacher M, Bui NK, Hood RD, Carl MA, Agnello DM, Schwarz S, Goodlett DR, Vollmer W, et al (2012) A widespread bacterial type VI secretion effector superfamily identified using a heuristic approach. Cell Host Microbe 11: 538–549

Saitou N & Nei M (1987) The neighbor-joining method: a new method for reconstructing phylogenetic trees. Mol Biol Evol 4: 406–425

Salomon D, Gonzalez H, Updegraff BL & Orth K (2013) Vibrio parahaemolyticus Type VI secretion system 1 Is activated in marine conditions to target bacteria, and is differentially regulated from system 2. PLoS One 8: e61086

Salomon D, Kinch LN, Trudgian DC, Guo X, Klimko JA, Grishin N V., Mirzaei H & Orth K (2014a) Marker for type VI secretion system effectors. Proc Natl Acad Sci 111: 9271–9276

Salomon D, Klimko JA & Orth K (2014b) H-NS regulates the Vibrio parahaemolyticus type VI secretion system 1. Microbiol (United Kingdom) 160: 1867–1873

Salomon D, Klimko JA, Trudgian DC, Kinch LN, Grishin N V., Mirzaei H & Orth K (2015) Type VI secretion system toxins horizontally shared between marine bacteria. PLoS Pathog 11: 1–20

Schindelin J, Arganda-Carreras I, Frise E, Kaynig V, Longair M, Pietzsch T, Preibisch S, Rueden C, Saalfeld S, Schmid B, et al (2012) Fiji: an open-source platform for biological-image analysis. Nat Methods 9: 676–682

Shneider MM, Buth SA, Ho BT, Basler M, Mekalanos JJ & Leiman PG (2013) PAAR-repeat proteins sharpen and diversify the type VI secretion system spike. Nature 500: 350–353

Silverman JM, Agnello DM, Zheng H, Andrews BT, Li M, Catalano CE, Gonen T & Mougous JD (2013) Haemolysin coregulated protein is an exported receptor and chaperone of type VI secretion substrates. Mol Cell 51: 584–593

Speare L, Cecere AG, Guckes KR, Smith S, Wollenberg MS, Mandel MJ, Miyashiro T & Septer AN (2018) Bacterial symbionts use a type VI secretion system to eliminate competitors in their natural host. Proc Natl Acad Sci U S A 115: E8528–E8537

Tran L, Nunan L, Redman R, Mohney L, Pantoja C, Fitzsimmons K & Lightner D (2013) Determination of the infectious nature of the agent of acute hepatopancreatic necrosis syndrome affecting penaeid shrimp. Dis Aquat Organ 105: 45–55

Unterweger D, Kostiuk B, Ötjengerdes R, Wilton A, Diaz-Satizabal L & Pukatzki S (2015) Chimeric adaptor proteins translocate diverse type VI secretion system effectors in Vibrio cholerae. EMBO J 34: 2198–210

Wang J, Brackmann M, Castaño-Díez D, Kudryashev M, Goldie KN, Maier T, Stahlberg H & Basler M (2017) Cryo-EM structure of the extended type VI secretion system sheath-tube complex. Nat Microbiol 2: 1507–1512

Wettstadt S, Wood TE, Fecht S & Filloux A (2019) Delivery of the Pseudomonas aeruginosa phospholipase effectors PldA and PldB in a VgrG- And H2-T6SS-dependent manner. Front Microbiol 10: 1718

Yu Y, Yang H, Li J, Zhang P, Wu B, Zhu B, Zhang Y & Fang W (2012) Putative type VI secretion systems of Vibrio parahaemolyticus contribute to adhesion to cultured cell monolayers. Arch Microbiol 194: 827–835

Zhang L & Orth K (2013) Virulence determinants for Vibrio parahaemolyticus infection. Curr Opin Microbiol 16: 70–77

