## Supplemental Data for "A binary effector module secreted by a type VI secretion system"

**Supplementary Figures S1-S5**

**Supplementary Movies S1-S2**

**Supplementary Tables S1-S3**

**Supplementary Datasets S1-S3**

**Supplementary References**

### Supplementary Figures

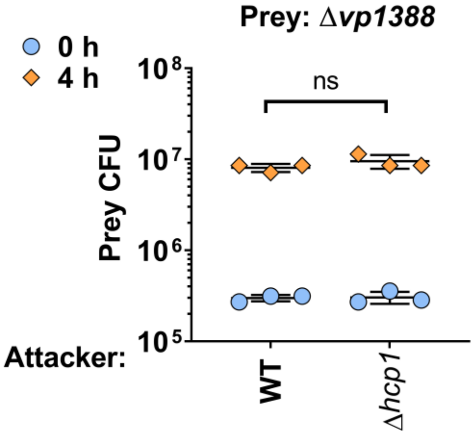


**Supplementary Figure S1. Deletion of *vp1388* does not affect bacterial immunity against T6SS1-mediated attacks.** Viability counts of *V. parahaemolyticus* Δ*vp1388* prey before (0 h) and after (4 h) co-incubation with the indicated *V. parahaemolyticus* attackers on media containing 3% NaCl at 30°C. Prey contain an empty plasmid providing a selection marker. Data are shown as the mean ± SD. Statistical significance between samples at the 4 h timepoint by an unpaired, two-tailed Student's *t*-test is denoted above. A significant difference was considered as $P < 0.05$. ns, no significant difference. Δ*hcp1* was used as a $T6SS1^-$ control strain.

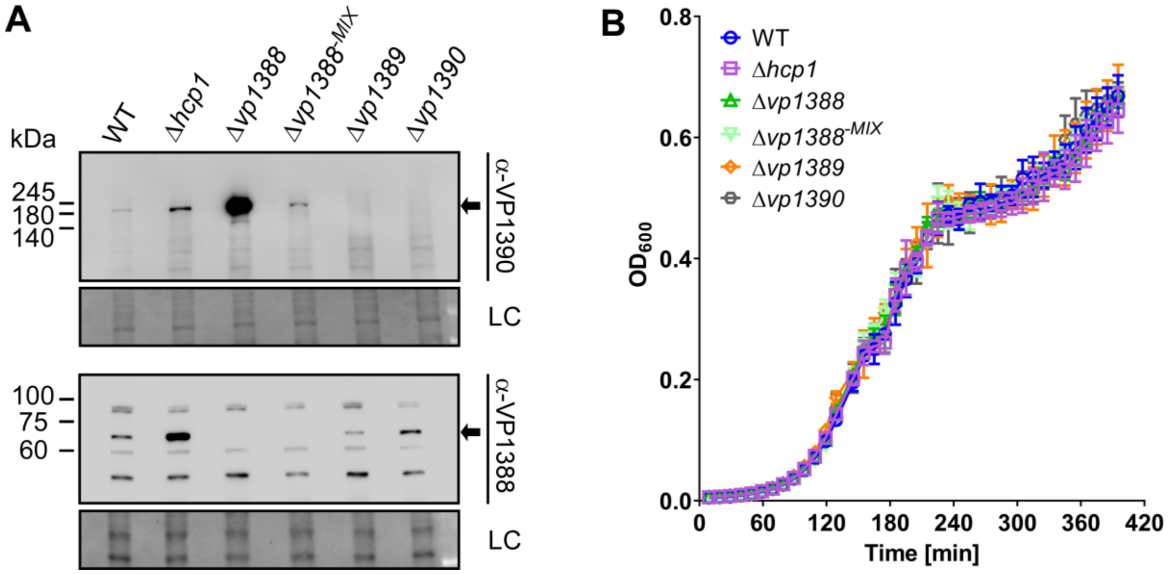


**Supplementary Figure S2. Generating a *vp1388* mutant with no polar effect on VP1390 expression. A)** Expression of VP1390 and VP1388 from *V. parahaemolyticus* mutant strains. Samples were treated with 20 µM phenamil to activate surface sensing in media containing 3% NaCl at 30°C for 5 h. Loading control (LC) is shown for total protein lysates. Arrows denote bands corresponding to VP1390 and VP1388. **B)** The growth of *V. parahaemolyticus* mutants in media containing 3% NaCl at 30°C, as measured by $OD_{600}$ readings. Data are shown as the mean ± SD. n = 4. WT, wild-type.

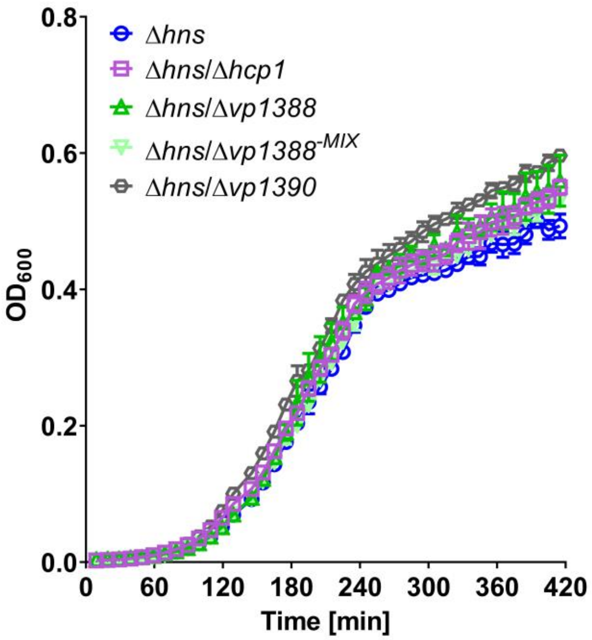


**Supplementary Figure S3. Mutants in *vp1388* and *vp1390* do not affect bacterial growth in a Δ*hns* background.** The growth of *V. parahaemolyticus* mutants in media containing 3% NaCl at 30°C, as measured by $OD_{600}$ readings. Data are shown as the mean ± SD. n = 4. WT, wild-type.

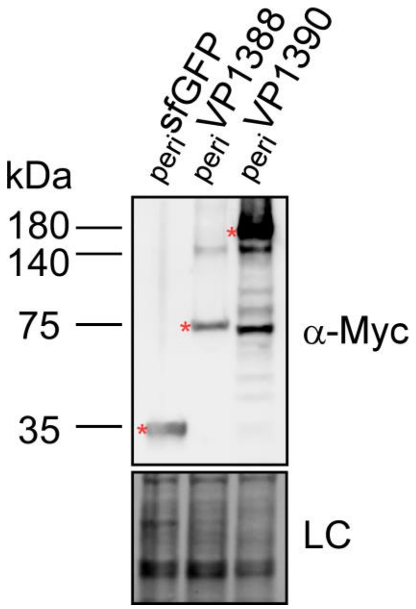


**Supplementary Figure S4. Expression of periplasm-targeted VP1388 and VP1390.** The expression of C-terminal Myc-tagged sfGFP, VP1388, and VP1390 fused to an N-terminal PelB signal peptide ($_{peri}$sfGFP, $_{peri}$VP1388, and $_{peri}$VP1390, respectively) was detected upon expression from an arabinose-inducible vector in *E. coli*, using anti-Myc antibodies. Bands corresponding to the protein expressed at the expected size are denoted by red asterisks. Loading control (LC) is shown for total protein lysates.

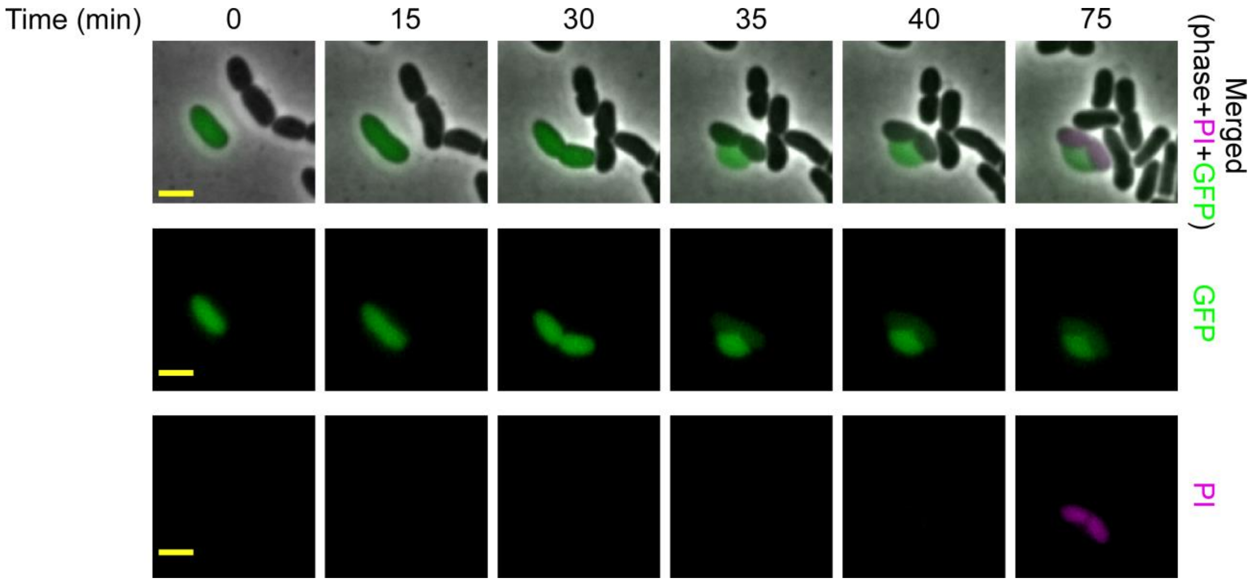


**Supplementary Figure S5. Lysis of sensitive prey cells is preceded by a bleb originating from the septum.** Representative image from time-lapse microscopy of competition between *V. parahaemolyticus* ∆*hns* (T6SS$^+$) or ∆*hns*/∆*hcp1* (T6SS$^-$) attackers and *V. parahaemolyticus* ∆*vp1389* prey that express GFP, as shown in Figure 5. Attacker and prey were mixed (2:1 ratio) and spotted on LB agarose pads supplemented with propidium iodide (PI; pink). Merging of the phase contrast, GFP (green) and PI (pink) channels, as well as the PI or GFP channels alone are shown. Scale bar = 2 µm.

### Supplementary Movies

**Supplementary Movie S1. Time-lapse microscopy of *E. coli* expressing periplasm-targeted VP1390, VP1388, or sfGFP.** *E. coli* expressing arabinose-inducible VP1390, VP1388, or sfGFP fused to an N-terminal PelB signal peptide were spotted on LB agarose pads supplemented with kanamycin (to maintain the plasmid), 0.2% arabinose, and propidium iodide (PI; pink). Cells were imaged every 5 minutes for 4 hours. Bar = 5 µm.

**Supplementary Movie S2. Time-lapse microscopy of Δ*vp1389* prey cells during competition**. *V. parahaemolyticus* Δ*vp1389* prey cells expressing sfGFP from a plasmid were visualized using a fluorescent microscope upon co-incubation with *V. parahaemolyticus* Δ*hns* ($T6SS1^+$) or Δ*hns*/Δ*hcp1* ($T6SS1^-$) attackers under T6SS1-inducing conditions on LB agarose pads supplemented with propidium iodide (PI; pink). Cells were imaged every 4 minutes for 1 hour. Bar = 5 µm.

### Supplementary Tables

#### Supplementary Table S1. A list of bacterial strains used in this work.

| Strain name | Genotype | Comments | Source |
|---|---|---|---|
| *Vibrio parahaemolyticus* RIMD 2210633 | Wild type | Used for generating deletion strains, as an attacker in bacterial competition assays, and as a template for PCR amplification. | Obtained from Kim Orth; (Makino *et al*, 2003) |
| *Vibrio parahaemolyticus* RIMD 2210633 Δ*hcp1* | Δ*hcp1* | RIMD 2210633 derivative containing an in-frame deletion of *vp1393;* used as an attacker in bacterial competition assays and for VP1388 and VP1390 expression. | This study |
| *Vibrio parahaemolyticus* RIMD 2210633 Δ*vp1388* | Δ*vp1388* | RIMD 2210633 derivative containing an in-frame deletion of *vp1388*; used as an attacker and prey in bacterial competition assays. | This study |
| *Vibrio parahaemolyticus* RIMD 2210633 Δ*vp1388*$^{-MIX}$ | Δ*vp1388*$^{-MIX}$ | RIMD 2210633 derivative containing a deletion of the *vp1388* region encoding a MIX domain (corresponding to amino acids 242-423); used as an attacker in bacterial competition assays. | This study |
| *Vibrio parahaemolyticus* RIMD 2210633 Δ*vp1389* | Δ*vp1389* | RIMD 2210633 derivative containing an in-frame deletion of *vp1389*; used as an attacker and prey in bacterial competition assays, and in immunoprecipitation assays. | This study |
| *Vibrio parahaemolyticus* RIMD 2210633 Δ*vp1390* | Δ*vp1390* | RIMD 2210633 derivative containing an in-frame deletion of *vp1390*; used as an attacker and prey in bacterial competition assays. | This study |
| *Vibrio parahaemolyticus* RIMD 2210633 Δ*hns* | Δ*hns* | RIMD 2210633 derivative containing an in-frame deletion of *vp1133*; used as an attacker and prey in | This study |

| | | | |
|---|---|---|---|
| | | bacterial competition assays, and in secretion assays. | |
| *Vibrio parahaemolyticus* RIMD 2210633 Δ*hns*/Δ*hcp1* | Δ*hns*/Δ*hcp1* | RIMD 2210633 derivative containing an in-frame deletion of *vp1133* and of *vp1393*; used as an attacker in bacterial competition assays, in secretion assays, and in immunoprecipitation assays. | This study |
| *Vibrio parahaemolyticus* RIMD 2210633 Δ*hns*/Δ*vp1388*$^{-MIX}$ | Δ*hns*/Δ*vp1388*$^{-MIX}$ | RIMD 2210633 derivative containing an in-frame deletion of *vp1133* and of the *vp1388* region encoding a MIX domain (corresponding to amino acids 242-423); used as an attacker in bacterial competition assays, and in secretion assays. | This study |
| *Vibrio parahaemolyticus* RIMD 2210633 Δ*hns*/Δ*vp1390* | Δ*hns*/Δ*vp1390* | RIMD 2210633 derivative containing an in-frame deletion of *vp1133* and of *vp1390*; used as an attacker in bacterial competition assays, and in secretion assays. | This study |
| *Vibrio parahaemolyticus* RIMD 2210633 Δ*hns*/Δ*hcp1*/Δ*vp1390* | Δ*hns*/Δ*hcp1*/Δ*vp1390* | RIMD 2210633 derivative containing an in-frame deletion of *vp1133*, *vp1393*, and *vp1390*; used for immunoprecipitation assays. | This study |
| *Escherichia coli* DH5α (λ pir) | K-12 derivative laboratory strain containing λ pir | Used for plasmid maintenance and cloning | Obtained from Eric V. Stabb |
| *Escherichia coli* XL1 Blue | Laboratory strain | Used as prey in bacterial competition assays | Purchased from Addgene |
| *Escherichia coli* sAJM.1506 | MG1655-derivative; “Marionette” strain | Used for protein expression, toxicity assays, and microscopy | Purchased from Addgene; (Meyer *et al*, 2019) |
| *Escherichia coli* BL21 (DE3) | Laboratory strain | Used for protein expression | Obtained from Kim Orth |

**Supplementary Table S2. A list of plasmids used in this work.**

| Plasmid name | Description | Comments | Source |
|---|---|---|---|
| pDM4 | a $Cm^R$ and oriVR6K-containing suicide vector | Used to generate deletions in *Vibrio* genomes | (O'Toole *et al*, 1996) |
| pDM4:*vp1388* | pDM4 containing 1 kb upstream and 1 kb downstream of *vp1388* in its MCS | Used to delete *vp1388* in *V. parahaemolyticus* RIMD 2210633 | (Salomon *et al*, 2014a) |
| pDM4:*vp1388*$^{-MIX}$ | pDM4 containing 1 kb upstream and 1 kb downstream of the *vp1388* region encoding the MIX domain in its MCS | Used to delete the *vp1388* region encoding the MIX domain (corresponding to amino acids 242-423) in *V. parahaemolyticus* RIMD 2210633 | This study |
| pDM4:*vp1389* | pDM4 containing 1 kb upstream and 1 kb downstream of *vp1389* in its MCS | Used to delete *vp1389* in *V. parahaemolyticus* RIMD 2210633 | This study |
| pDM4:*vp1390* | pDM4 containing 1 kb upstream and 1 kb downstream of *vp1390* in its MCS | Used to delete *vp1390* in *V. parahaemolyticus* RIMD 2210633 | This study |
| pDM4:*hns* | pDM4 containing 1 kb upstream and 1 kb downstream of *vp1133* in its MCS | Used to delete *hns* in *V. parahaemolyticus* RIMD 2210633 | (Salomon *et al*, 2014b) |
| pDM4:*hcp1* | pDM4 containing 1 kb downstream and 1 kb upstream of *vp1393* in its MCS | Used to delete *vp1393* in *V. parahaemolyticus* RIMD 2210633 | (Salomon *et al*, 2013) |
| $pBAD^K$/Myc-His | pBR322 ori-containing plasmid harboring a $Kan^R$ cassette, araC, and an MCS following a P*bad* promoter | Used for arabinose-inducible expression | (Salomon *et al*, 2013) |
| pPER5 | $pBAD^K$/Myc-His with a PelB signal peptide inserted at the 5' end of the MCS | Used for arabinose-inducible expression of proteins targeted to the periplasm in *E. coli* | (Dar *et al*, 2018) |
| pBAD33.1 | p15A ori-containing plasmid carrying a $Cm^R$ gene, araC, and an MCS following a P*bad* promoter. | Used as template to construct $pBAD33.1^F$ | Purchased from Addgene; (Chung & Raetz, 2010) |
| $pBAD33.1^F$ | pBAD33.1 with a FLAG tag inserted at the 3' end of the MCS | Used for arabinose-inducible expression | (Fridman *et al*, 2020) |
| $pVP1388^{peri}$ | pPER5 plasmid containing the CDS of *vp1388* in frame with the N-terminal PelB signal peptide and the | Used for arabinose-inducible expression of periplasm-targeted VP1388 in *E. coli* | This study |

| | | | |
|---|---|---|---|
| | C-terminal Myc-His tag | | |
| pVP1388$^{Myc}$ | pBAD$^{K}$/Myc-His plasmid containing the CDS of *vp1388* in frame with a C-terminal Myc-His tag | Used for arabinose-inducible expression of VP1388 in *E. coli* | This study |
| pVP1388 | pBAD33.1$^{F}$ plasmid containing the CDS of *vp1388* in frame with the C-terminal FLAG tag | Used for arabinose-inducible expression of VP1388 in *Vibrio* | This study |
| pVP1389 | pBAD/Myc-His containing the CDS of *vp1389* in frame with the C-terminal Myc-His tag | Used for arabinose-inducible expression of VP1389 in *Vibrio* | This study |
| pVP1390$^{peri}$ | pPER5 plasmid containing the CDS of *vp1390* in frame with the N-terminal PelB signal peptide and the C-terminal Myc-His tag | Used for arabinose-inducible expression of periplasm-targeted VP1390 in *E. coli* | This study |
| pVP1390 | pBAD33.1 plasmid containing the CDS of *vp1390* in frame with the C-terminal FLAG tag | Used for arabinose-inducible expression of VP1390 in *Vibrio* and in *E. coli* | This study |
| pPoNi | pBAD33.1 plasmid containing the CDS of *b5c30_rs14460* from *V. parahaemolyticus* 12-297/B in frame with the C-terminal FLAG tag | Used for arabinose-inducible expression of PoNi in *E. coli* | (Jana *et al*, 2019) |
| pPoNe$^{D335A}$ | pBAD/Myc-His containing the CDS of *b5c30_rs14465* from *V. parahaemolyticus* 12-297/B carrying a mutation in the active site D335A, in frame with the C-terminal Myc-His tag | Used for arabinose-inducible expression of PoNe$^{D335A}$ in *E. coli* | (Jana *et al*, 2019) |
| psfGFP$^{peri}$ | pPER5 plasmid containing the CDS of sfGFP in frame with the N-terminal PelB signal peptide and the C-terminal Myc-His tag | Used for arabinose-inducible expression of sfGFP in *E. coli* | This study |
| psfGFP | pBAD33.1$^{F}$ plasmid containing the CDS of sfGFP in frame with | Used for arabinose-inducible expression of sfGFP in *Vibrio* | This study |

| | | | |
|---|---|---|---|
| | the C-terminal FLAG tag | | |
| pBC3020 | pBAD33.1$^{F}$ plasmid containing the CDS of BC3020 in frame with the C-terminal FLAG tag | Used for arabinose-inducible expression of BC3020 in *Vibrio* | (Jana *et al*, 2019) |
| pGFP | a Spec$^{R}$-containing, high copy number plasmid for constitutive expression of GFP in vibrios | Used for constitutive expression of GFP in *V. parahaemolyticus* prey visualized under the microscope | (Ritchie *et al*, 2012) |

**Supplementary Table S3. A list of primers used in this work.**

| Primer name | Sequence (5' to 3') | Description |
|---|---|---|
| VP1388_MIX_SacI_UP_F | CACCGAGCTCCCTCAACCTTCGGCTATCAGT | Used to amplify 1 kb upstream of the *vp1388* MIX domain-encoding region to construct pDM4:*vp1388*$^{-MIX}$ |
| VP1388_MIX_UP_R | CCGGGATCACTTTCCAACACGTTATCCCACTCATCC | |
| VP1388_MIX_DN_F | GGATAACGTGTTGGAAAGTGATCCCGGAACCAAG | Used to amplify 1 kb downstream of the *vp1388* MIX domain-encoding region to construct pDM4:*vp1388*$^{-MIX}$ |
| VP1388_MIX_SalI_DN_R | CCACGTCGACCCTAGTTCAATATACTCACCCG | |
| VP1389_SacI_UP_F | CACCGAGCTCAATTACGGAAAATGGTTAC | Used to amplify 1 kb upstream of *vp1389* to construct pDM4:*vp1389* |
| VP1389_BamHI_UP_R | CAACGGATCCCTTATATTCTCTTCGTTATC | |
| VP1389_BamHI_DN_F | CAACGGATCCGCCTTGTAGAGAACTGCAATG | Used to amplify 1 kb downstream of *vp1389* to construct pDM4:*vp1389* |
| VP1389_SalI_DN_R | CAACGTCGACGCAAATTTCCCTAACGGAG | |
| VP1390_SphI_UP_F | CACCGCATGCAATGCTCAACAGTGGGAAG | Used to amplify 1 kb upstream of *vp1390* to construct pDM4:*vp1390* |
| VP1390_BamHI_UP_R | CAACGGATCCAAATCCCCCTCTTTAACAAAATC | |
| VP1390_BamHI_DN_F | CACCGGATCCATGCAAAGAAAAACCGCC | Used to amplify 1 kb downstream of *vp1390* to construct pDM4:*vp1390* |
| VP1390_SpeI_DN_R | CAACACTAGTTGGTGAAACGGGCGCTGG | |
| pVP1388$^{peri}$_F | CCCAGCCGGCGATGGCCATGTGGAGAATTCAAATGCATTTAG | Used to amplify *vp1388* to construct |
| pVP1388$^{peri}$_R | TTTTGTTCGGGCCCAAGCTTACTTGGCTGATCTTTTTTCTTATAAG | |

| | | |
|---|---|---|
| | | pVP1388$^{\text{peri}}$ |
| pVP1388$^{\text{Myc}}$_F | GCTAACAGGAGGAATTAACCTTGTGGAGAATTCAAATGC | Used to amplify *vp1388* to construct pVP1388$^{\text{Myc}}$ |
| pVP1388$^{\text{Myc}}$_R | TTTTGTTCGGGCCCAAGCTTACTTGGCTGATCTTTTTTCTTATAAGCC | |
| pVP1388_F | CTTTAAGAAGGAGATATACATTTGTGGAGAATTCAAATGC | Used to amplify *vp1388* to construct pVP1388 |
| pVP1388_R | CGTCGTCATCCTTGTAATCACTTGGCTGATCTTTTTTCTTATAAGC | |
| pVP1389_F | GCTAACAGGAGGAATTAACCATGAATCTACGTCATACATTATGC | Used to amplify *vp1389* to construct pVP1389 |
| pVP1389_R | TTTTGTTCGGGCCCAAGCTTTAAATCCCCCTCTTTAAC | |
| pVP1390$^{\text{peri}}$_F | CCCAGCCGGCGATGGCCATGAGCCTTGTAGAGAACTGCAATG | Used to amplify *vp1390* to construct pVP1390$^{\text{peri}}$ |
| pVP1390$^{\text{peri}}$_R | TTTTGTTCGGGCCCAAGCTTGGAAGGGTTATCAAGTGACTTTTC | |
| pVP1390_F | CTTTAAGAAGGAGATATACATATGAGCCTTGTAGAGAACTGC | Used to amplify *vp1390* to construct pVP1390 |
| pVP1390_R | CGTCGTCATCCTTGTAATCGGAAGGGTTATCAAGTGACTTTTC | |
| psfGFP$^{\text{peri}}$_F | CCCAGCCGGCGATGGCCATGGTGAGCAAGGGCGAGGAG | Used to amplify *sfgfp* to construct psfGFP$^{\text{peri}}$ |
| psfGFP$^{\text{peri}}$_R | TTTTGTTCGGGCCCAAGCTTCTTGTACAGCTCGTCCATGCC | |
| psfGFP_F | CTTTAAGAAGGAGATATACATATGGTGAGCAAGGGCGAGGAGC | Used to amplify *sfgfp* to construct psfGFP |
| psfGFP_R | CGTCGTCATCCTTGTAATCCTTGTACAGCTCGTCCATGCCG | |

### Supplementary Datasets

**Supplementary Dataset S1 - Bacterial genomes containing *vp1388-90* homologous operons**

**Supplementary Dataset S2 - Summary of homologous operons identified in bacterial genomes**

**Supplementary Dataset S3 - Proximity of homologous operons to T6SS core components**

### Supplementary References


Chung HS & Raetz CRH (2010) Interchangeable domains in the Kdo transferases of escherichia coli and haemophilus influenzae. *Biochemistry* 49: 4126–4137

Dar Y, Salomon D & Bosis E (2018) The antibacterial and anti-eukaryotic Type VI secretion system MIX-effector repertoire in Vibrionaceae. *Mar Drugs* 16: 433

Fridman CM, Keppel K, Gerlic M, Bosis E & Salomon D (2020) A comparative genomics methodology reveals a widespread family of membrane-disrupting T6SS effectors. *Nat Commun* 11: 1085

Jana B, Fridman CM, Bosis E & Salomon D (2019) A modular effector with a DNase domain and a marker for T6SS substrates. *Nat Commun* 10: 3595

Makino K, Oshima K, Kurokawa K, Yokoyama K, Uda T, Tagomori K, Iijima Y, Najima M, Nakano M, Yamashita A, *et al* (2003) Genome sequence of Vibrio parahaemolyticus: a pathogenic mechanism distinct from that of V cholerae. *Lancet* 361: 743–749

Meyer AJ, Segall-Shapiro TH, Glassey E, Zhang J & Voigt CA (2019) Escherichia coli “Marionette” strains with 12 highly optimized small-molecule sensors. *Nat Chem Biol* 15: 196–204

O’Toole R, Milton DL & Wolf-Watz H (1996) Chemotactic motility is required for invasion of the host by the fish pathogen Vibrio anguillarum. *Mol Microbiol* 19: 625–637

Ritchie JM, Rui H, Zhou X, Iida T, Kodoma T, Ito S, Davis BM, Bronson RT & Waldor MK (2012) Inflammation and Disintegration of Intestinal Villi in an Experimental Model for Vibrio parahaemolyticus-Induced Diarrhea. *PLoS Pathog* 8: e1002593

Salomon D, Gonzalez H, Updegraff BL & Orth K (2013) Vibrio parahaemolyticus Type VI secretion system 1 Is activated in marine conditions to target bacteria, and is differentially regulated from system 2. *PLoS One* 8: e61086

Salomon D, Kinch LN, Trudgian DC, Guo X, Klimko JA, Grishin N V., Mirzaei H & Orth K (2014a) Marker for type VI secretion system effectors. *Proc Natl Acad Sci* 111: 9271–9276

Salomon D, Klimko JA & Orth K (2014b) H-NS regulates the Vibrio parahaemolyticus type VI secretion system 1. *Microbiol (United Kingdom)* 160: 1867–1873
